# Dynamics in Fip1 regulate eukaryotic mRNA 3'-end processing

**DOI:** 10.1101/2021.07.07.451483

**Authors:** Ananthanarayanan Kumar, Conny W.H. Yu, Juan B. Rodríguez-Molina, Xiao-Han Li, Stefan MV Freund, Lori A Passmore

**Affiliations:** MRC Laboratory of Molecular Biology, Cambridge UK; Department of Molecular, Cellular, and Developmental Biology, Yale University, New Haven, CT 06511, USA

## Abstract

Cleavage and polyadenylation factor (CPF/CPSF) is a multiprotein complex essential for mRNA 3’-end processing in eukaryotes. It contains an endonuclease that cleaves pre-mRNAs, and a polymerase that adds a poly(A) tail onto the cleaved 3’-end. Several CPF subunits, including Fip1, contain intrinsically-disordered regions (IDRs). IDRs within multiprotein complexes can be flexible, or can become ordered upon interaction with binding partners. Here, we show that yeast Fip1 anchors the poly(A) polymerase Pap1 onto CPF via an interaction with zinc finger 4 of another CPF subunit, Yth1. We also reconstitute a fully recombinant 850-kDa CPF. By incorporating selectively-labelled Fip1 into recombinant CPF, we could study the dynamics of this single protein within the megadalton complex using nuclear magnetic resonance spectroscopy (NMR). This reveals that a Fip1 IDR that connects the Yth1- and Pap1-binding sites remains highly dynamic within CPF. Together, our data suggest that Fip1 dynamics mediate conformational transitions within the 3’-end processing machinery to coordinate cleavage and polyadenylation.

## Introduction

Protein-coding genes in eukaryotes are transcribed by RNA polymerase II (Pol II) into precursor messenger RNAs (pre-mRNAs). Pre-mRNAs are modified by the addition of a 7-methylguanosine cap at the 5’-end, splicing, and 3’-end processing (Hocine et al. 2010). The 3’-end of an mRNA is formed by a two-step reaction involving endonucleolytic cleavage at a specific site and the addition of a stretch of polyadenosines (a poly(A) tail) to the new free 3’ hydroxyl (Zhao et al. 1999). Poly(A) tails are essential for export of mature mRNAs into the cytoplasm, for their subsequent translation into proteins, and in determining mRNA half-life. Defects in 3’-end processing are associated with human diseases including cancer, β-thalassemia and spinal muscular atrophy (Curinha et al. 2014). Understanding the mechanistic basis of how poly(A) tails are added to mRNA 3’-ends and how polyadenylation is coordinated with other mRNA processing steps is therefore of great importance.

Eukaryotic 3’-end processing is carried out by a set of conserved multiprotein complexes that includes the cleavage and polyadenylation factor (CPF in yeast or CPSF in humans) and accessory cleavage factors (CF IA and CF IB in yeast, or CF Im, CF IIm and CstF in humans) (Kumar et al. 2019). In yeast, CPF is comprised of three enzymatic modules: a five-subunit polymerase module containing the poly(A) polymerase Pap1, a three-subunit nuclease module containing the endonuclease Ysh1, and a six-subunit phosphatase module that includes two protein phosphatases (Glc7 and Swd2) that regulate transcription (Casanal et al. 2017). Most CPF subunits are conserved across all eukaryotes.

Insights into the molecular basis of polyadenylation have been obtained through structural and biochemical studies. For example, a crystal structure of Pap1 in complex with ATP, poly(A) RNA and Mg^2+^ confirmed that a two-metal-ion dependent nucleotidyl transfer mechanism is employed in poly(A) tail synthesis (Balbo and Bohm 2007). Together with kinetic studies, this structure provided a molecular basis for nucleotide specificity. Pap1 is assembled into the polymerase module along with the scaffolding proteins Cft1 and Pfs2, the zinc finger-containing protein Yth1, and the low sequence complexity protein Fip1 (Casanal et al. 2017). A similar mammalian polymerase module (mPSF) is sufficient for specific and efficient mRNA polyadenylation *in vitro* (Schonemann et al. 2014). Cryo-electron microscopy (cryoEM) structures of the polymerase modules from yeast and human revealed an extensive network of interactions between beta propeller domains of Pfs2 and Cft1 (WDR33 and CPSF160 in human) (Casanal et al. 2017; Clerici et al. 2017; Sun et al. 2018). The structures also provided a rationale for how WDR33 and CPSF30 (Yth1 in yeast) bind specific sequences in RNA (Clerici et al. 2018; Sun et al. 2018).

Fip1 and Pap1 interact directly with each other and a crystal structure of yeast Pap1 bound to residues 80–105 of Fip1 provided the molecular details of their interaction (Meinke et al. 2008). Fip1 has also been reported to interact with other CPF and CF IA components such as Pta1, Yth1 and Rna14 (Preker et al. 1995; Barabino et al. 2000; Ohnacker et al. 2000; Tacahashi et al. 2003; Ghazy et al. 2009; Casanal et al. 2017) but neither Fip1 nor Pap1 were visible in cryoEM studies. Fip1 has an N-terminal acidic stretch followed by a Pro-rich region (Preker et al. 1995; Kaufmann et al. 2004). This overall domain architecture of Fip1 is conserved, but human FIP1 (hFIP1) additionally contains C-terminal Arg-rich and Arg/Asp-rich domains compared to the yeast ortholog (Kaufmann et al. 2004). Biochemical and genetic experiments led to the hypothesis that Fip1 is an unstructured protein that acts as a flexible linker between Pap1 and CPF (Meinke et al. 2008; Ezeokonkwo et al. 2011). However, Fip1 structure has only been studied in isolation; whether Fip1 remains dynamic in the context of CPF remains unclear.

The nuclease module subunits are flexibly positioned with respect to the polymerase module (Hill et al. 2019; Zhang et al. 2020), but they are hypothesized to become fixed upon CPF activation (Sun et al. 2020). It is likely that a series of conformational transitions are required for CPF function: first the nuclease must be activated; second, the nuclease must be inactivated after cleavage has occurred; and finally, the RNA is transferred to Pap1’s active site to allow the poly(A) tail to be synthesized to the correct length. Conformational changes are frequently associated with dynamic IDRs (Fuxreiter et al. 2014; van der Lee et al. 2014); however, it remains unknown whether IDRs within CPF subunits mediate changes between states.

Here, we aimed to understand the function and dynamics of Fip1 using biochemical reconstitution, biophysical experiments and NMR spectroscopy. We find that isolated Fip1 is an intrinsically-disordered protein in solution, with defined binding sites for Yth1 and Pap1 that are connected by a low-complexity sequence. To fully characterize Fip1 as an essential component of the CPF, we reconstituted a recombinant 850-kDa CPF. This allowed us to incorporate an isotopically-labeled Fip1 into CPF for NMR studies, in which we show that Fip1 remains dynamic and largely disordered within CPF. Moreover, deletion of the highly flexible region in Fip1 impairs CPF nuclease activity. Together, our data reveal that Fip1 dynamics are important in regulating eukaryotic mRNA 3’-end processing.

## Results

### Yth1 zinc finger 4 binds Fip1 which in turn tethers Pap1 to the polymerase module

To investigate how Fip1 and Pap1 interact with other CPF subunits, we first studied the five-subunit polymerase module purified from a baculovirus-mediated insect cell overexpression system as previously described (Casanal et al. 2017). We used cryoEM to image this five-subunit complex (Supplemental Fig. S1). Selected 2D class averages show a central structure that resembles the Cft1-Pfs2-Yth1 scaffold identified previously in the four-subunit complex (Fig. 1A). In addition, a horseshoe-shaped structure was present in the 2D class averages at several different positions relative to Cft1-Pfs2-Yth1. This extra density resembles a 2D projection of the crystal structure of Pap1 (Bard et al. 2000) suggesting that Pap1 is positioned flexibly with respect to the Cft1-Pfs2-Yth1 scaffold. The lack of a defined position for Pap1 precluded high-resolution structure determination.

**Fig. 1.**
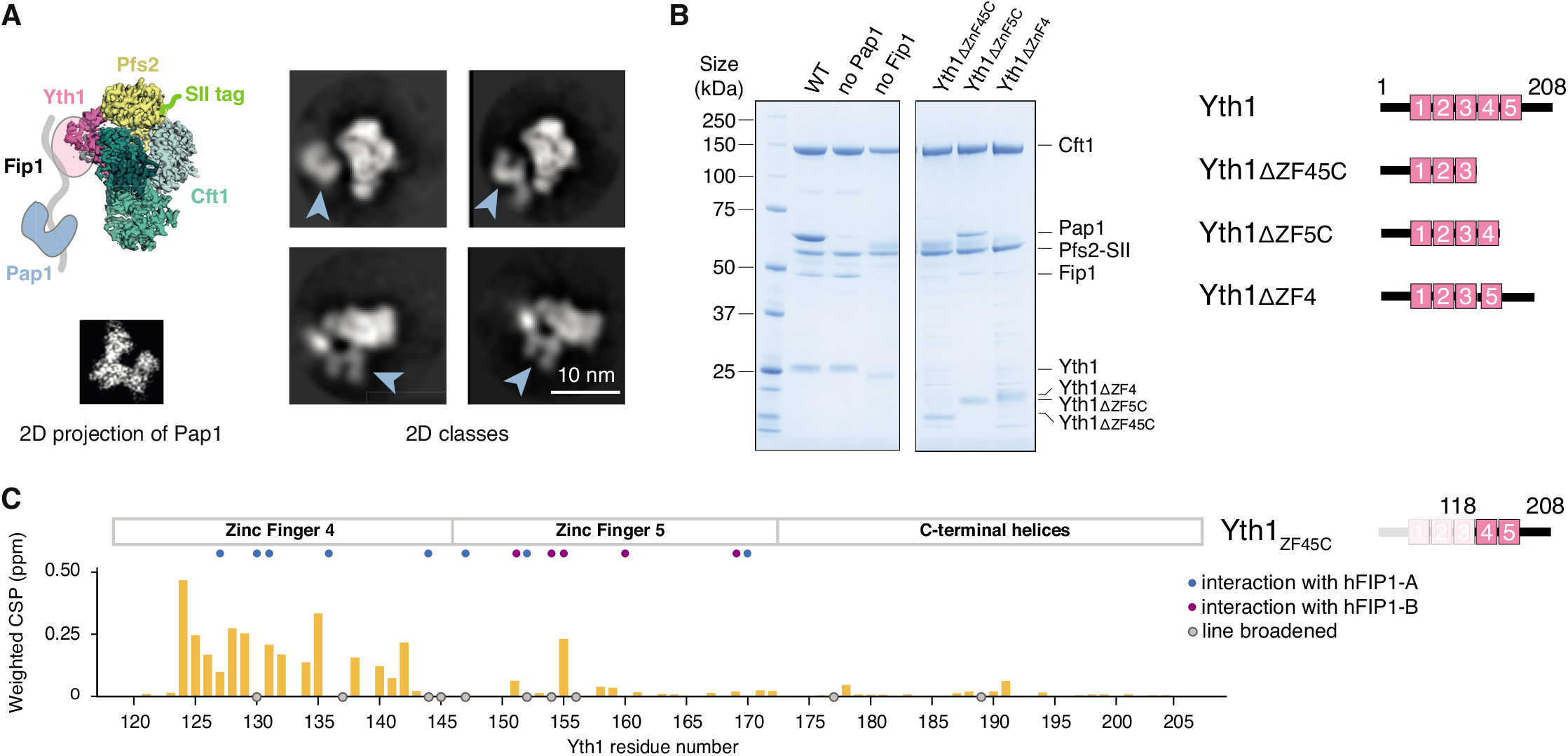
Fip1 binds Yth1 and tethers Pap1 to the polymerase module. **(A)** Cartoon representation (top left) and selected 2D class averages (right) from cryoEM of the polymerase module. The 2D averages show a central structure corresponding to the Cft1-Pfs2-Yth1 subunits, and an additional horseshoe-shaped density (blue arrowheads). A 2D projection of the crystal structure of Pap1 (PDB 3C66, bottom left) suggests that this horseshoe-shaped density is Pap1. SII, StrepII. **(B)** SDS-PAGE of pulldown assays using StrepII-tagged (SII) Pfs2 reveal interactions within the polymerase module. Domain diagrams of Yth1 indicate the constructs used (right). Fip1 is required for Pap1 interaction. When the interaction between Yth1 and Fip1 is compromised, Fip1 is not pulled down and therefore Pap1 is also absent. Yth1 subunits in the WT and ‘no Pap1’ complexes contain a His-tag, whereas Yth1 in the ‘no Fip1’ and Yth1 truncation complexes do not contain a His-tag. **(C)** Histogram showing chemical shift perturbations (CSP) in Yth1_ZF45C_ (residues 118–208) spectra upon Fip1_226_ binding. Grey circles indicate peaks that showed exchange broadening upon Fip1 binding. Most of the peaks that are perturbed are in zinc finger 4. Homologous residues contributing >50 Å^2^ buried surface area in the human FIP1–CPSF30 structure (Hamilton and Tong, 2020) are highlighted with blue (zinc finger 4) or purple (zinc finger 5) circles. In that structure, two hFIP1 molecules (hFIP1-A and hFIP1-B) are bound to one Yth1.

Next, to gain further insight into the architecture of the polymerase module, we investigated subunit interactions using pulldown assays. Using a StrepII tag on Pfs2, all five subunits were co-purified (Fig. 1B). When we removed Pap1 from the complex, the four remaining subunits were still associated. However, removal of Fip1 resulted in concomitant loss of Pap1 from the complex (Fig. 1B). Thus, Fip1 is essential for Pap1 association with the polymerase module. If Fip1 is dynamic, it may contribute to the variable positioning of Pap1 in EM images (Fig. 1A).

Fip1 is hypothesized to bind Yth1, which contains five zinc fingers. The N-terminal half of Yth1, including zinc fingers 1 and 2, interacts with Cft1 and Pfs2 (Casanal et al. 2017) while zinc fingers 2 and 3 interact with RNA (Clerici et al. 2018; Sun et al. 2018). The C-terminal half of Yth1 is not visible in cryoEM maps, suggesting that it may be flexible. To test whether the C-terminal region is required for interaction with other polymerase module subunits, we carried out pulldown assays with versions of Yth1 containing C-terminal deletions. Deletion of zinc finger 4 resulted in loss of Fip1 and Pap1 from the Pfs2 pulldowns (Fig. 1B). Deletion of zinc finger 5 and a C-terminal helical region reduced, but did not abolish, the association of Fip1 and Pap1. Deletion of regions spanning the zinc finger 4, zinc finger 5 and the C-terminal helical region resulted in a complete loss of Fip1 and Pap1 association with the complex. We therefore hypothesized that Yth1 zinc finger 4 is required for Fip1 incorporation into the fully recombinant polymerase module, in agreement with a previously proposed role for this zinc finger in Fip1 binding (Tacahashi et al. 2003). Zinc finger 5 may contribute more weakly to Fip1 binding.

To characterize the Yth1-Fip1 interaction, we performed NMR analysis of two Yth1 constructs: the entire C-terminal half (Yth1_ZF45C_; residues 118–208) or zinc finger 4 on its own (Yth1_ZF4_; residues 118–161) (Supplemental Fig. S2A-C). We assigned backbone resonances of Yth1_ZF45C_ and mapped chemical shift perturbations upon addition of a Fip1 construct containing residues 1–226 (Fip1_226_) (Fig. 1C, Supplemental Fig. S2D). (Details on the choice of Fip1_226_ construct are described in the next section). Addition of Fip1_226_ resulted in substantial chemical shift perturbations and line-broadening. The majority of changes (19/25 residues experiencing chemical shift perturbation or line-broadening) are located within residues in zinc finger 4, with a similar pattern in both Yth1_ZF45C_ and Yth1_ZF4_ spectra. The remaining changes are located in zinc finger 5. Thus, Yth1 zinc finger 4 is the primary interaction site for Fip1.

There are apparent domain boundaries on either side of Yth1 zinc finger 4 but this region contains very little secondary structure (Supplemental Fig. S2E-F). We confirmed a direct interaction of Yth1_ZF4_ and Fip1_226_ using isothermal calorimetry, with a measured binding affinity of 240 ± 40 nM (Supplemental Fig. S2G). We also found that zinc finger 4 in Yth1_ZF45C_ loses its structure upon incubation with EDTA, suggesting that the zinc ions are exposed and susceptible to metal chelation (Supplemental Fig. S2H). Interestingly, Yth1_ZF45C_ becomes resistant to loss of metal coordination after binding to Fip1_226_ as there is minimal change in the spectra in the presence of EDTA (Supplemental Fig. S2I). Taken together, Fip1 directly interacts with and stabilizes Yth1.

### Fip1 is largely disordered

Next, we examined the structure of Fip1. We used bioinformatics analyses to investigate the topology of the full-length, 327-amino acid protein (Fig. 2A). Overall, >75% of Fip1 (~250 residues) is predicted to be highly disordered including an N-terminal serine-rich acidic region (residues 1–60), a central low-complexity region (LCR) rich in serine and threonine (residues 110–180), and a C-terminal region with no net charge that is rich in asparagine, proline and phenylalanine (residues 243–327). This is consistent with previous evidence that isolated Fip1 is largely unfolded (Meinke et al. 2008; Ezeokonkwo et al. 2011). Interestingly, the only region that is predicted to have low disorder propensity (residues 193– 215) overlaps with sequences previously identified as important for Yth1 binding (residues 206–220) (Helmling et al. 2001) and is highly conserved across eukaryotes (Supplemental Fig. S3A).

**Fig. 2.**
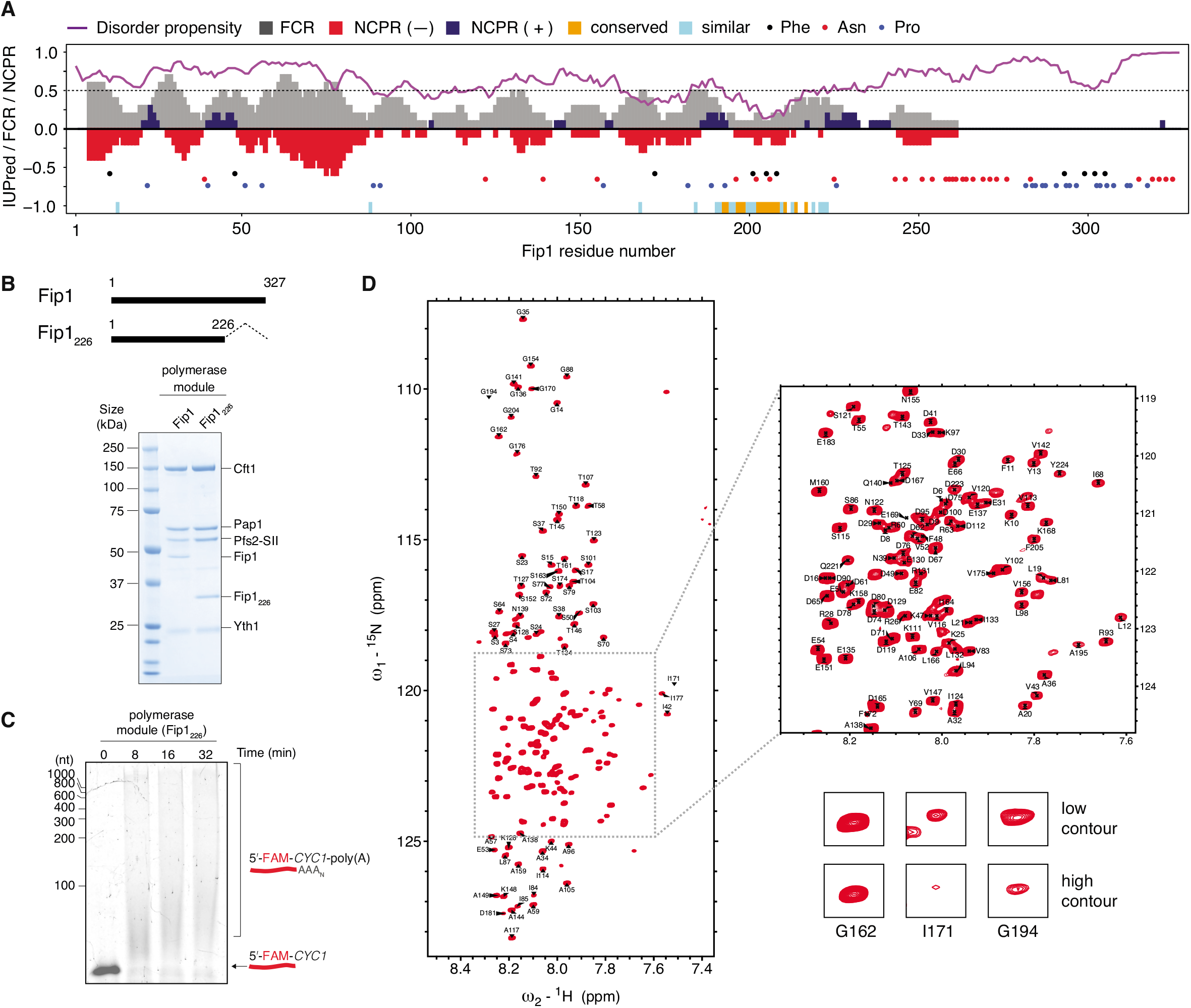
Fip1 residues 1–226 are sufficient for reconstitution of the polymerase module. **(A)** Bioinformatic analysis of Fip1 structure. Purple line indicates IUPRED2 disorder prediction score. Residues with an IUPred value over 0.5 (grey dotted line) are classified as having high propensity for being intrinsically disordered. Grey bars correspond to fraction of charged residues (FCR) over a sliding window of 10 residues. Red and blue bars represent net negative and positive charge per residue (NCPR) respectively, over the same window size. Sequence conservation is shown at the bottom of the plot (orange bars, conserved; cyan bars, similar). The colored circles highlight the distribution of phenylalanine (black), asparagine (red) and proline (blue) residues. **(B)** Schematic showing construct design of Fip1_226_ (residues 1–226; top) and SDS-PAGE of pulldown assays of polymerase module using StrepII-tagged (SII) Pfs2 (bottom). **(C)** *In vitro* polyadenylation assay with recombinant polymerase module containing Fip1_226_. A 42-nucleotide *CYC1* 3’-UTR with 5’-FAM label was used as a substrate in the polyadenylation assay and the reaction products were visualized on a denaturing urea polyacrylamide gel. This gel is representative of experiments performed two times. **(D)** ^1^H,^15^N 2D HSQC of Fip1_226_ shown with the assignment of backbone resonances. The inset highlights a crowded region of the spectra. Lower panels show examples of peaks. I171 and G194 show line broadening.

The C-terminal region is more poorly conserved than the N-terminal region, both in length and in amino acid composition (Supplemental Fig. S3B), suggesting that the C-terminus may not have a conserved functional role. A previous study showed that deletion of residues 220–327 of Fip1 had negligible effect on the viability of yeast cells and no significant effect on mRNA polyadenylation *in vitro* (Helmling et al. 2001). Asn-rich sequences are often aggregation prone. Since overexpression of isolated full-length Fip1 resulted in insoluble protein aggregates, we removed this sequence in a Fip1_226_ construct (Fip1 Δ227-327) that is stable for *in vitro* characterization. As a helical stretch is predicted close to the truncation point, we chose proline 226 as a natural helix breaker for the new C-terminus.

We co-expressed Fip1_226_ with the other polymerase module subunits in *Sf9* insect cells and performed pull-down assays using the StrepII-tag on Pfs2 (Fig. 2B). This showed that Fip1_226_ is incorporated into the recombinant polymerase module. In addition, this complex was active in *in vitro* polyadenylation assays (Fig. 2C). Together, this shows that Fip1_226_ is sufficient for *in vitro* reconstitution and polyadenylation activity of the polymerase module, confirming our prediction that residues beyond 226 on Fip1 are not essential for cleavage and polyadenylation.

Next, we analyzed Fip1_226_ using NMR. Several regions contain substantial signal attenuation due to line broadening, including residues between 170 and 220 (Supplemental Fig. S3C-E; Methods). We assigned 187 out of 226 (83%) backbone resonances (Fig. 2D). Although several regions have some propensity to form secondary structure (Supplemental Fig. S3E), Fip1_226_ generally has a narrow ^1^H dispersion in a ^1^H,^15^N 2D HSQC spectrum (Fig. 2D), consistent with Fip1 being a largely disordered protein.

### Fip1 contains independent binding sites for Yth1 and Pap1

Within multiprotein complexes, IDRs can be dynamic or they can become ordered upon binding other subunits (Fuxreiter et al. 2014; van der Lee et al. 2014). We next investigated the dynamics of Fip1 bound to other subunits. First, to identify the interaction sites for polymerase module subunits within Fip1, we made deletions in Fip1 and performed pulldown assays using the StrepII tag on Pfs2 (Fig. 3A). Residues 80–105 had previously been shown to mediate Fip1 interaction with Pap1 (Meinke et al. 2008) and therefore we did not assess that region in pulldown assays. We found that Fip1 residues 190–220 are required for Fip1 (and Pap1) interaction with the polymerase module. In contrast, deletion of the N-terminal acidic region (residues 1–60) or the central LCR (residues 110–180) had minimal effect on Fip1 interactions. These experiments therefore suggest that Fip1 residues 190–220 interact with Yth1.

**Fig. 3.**
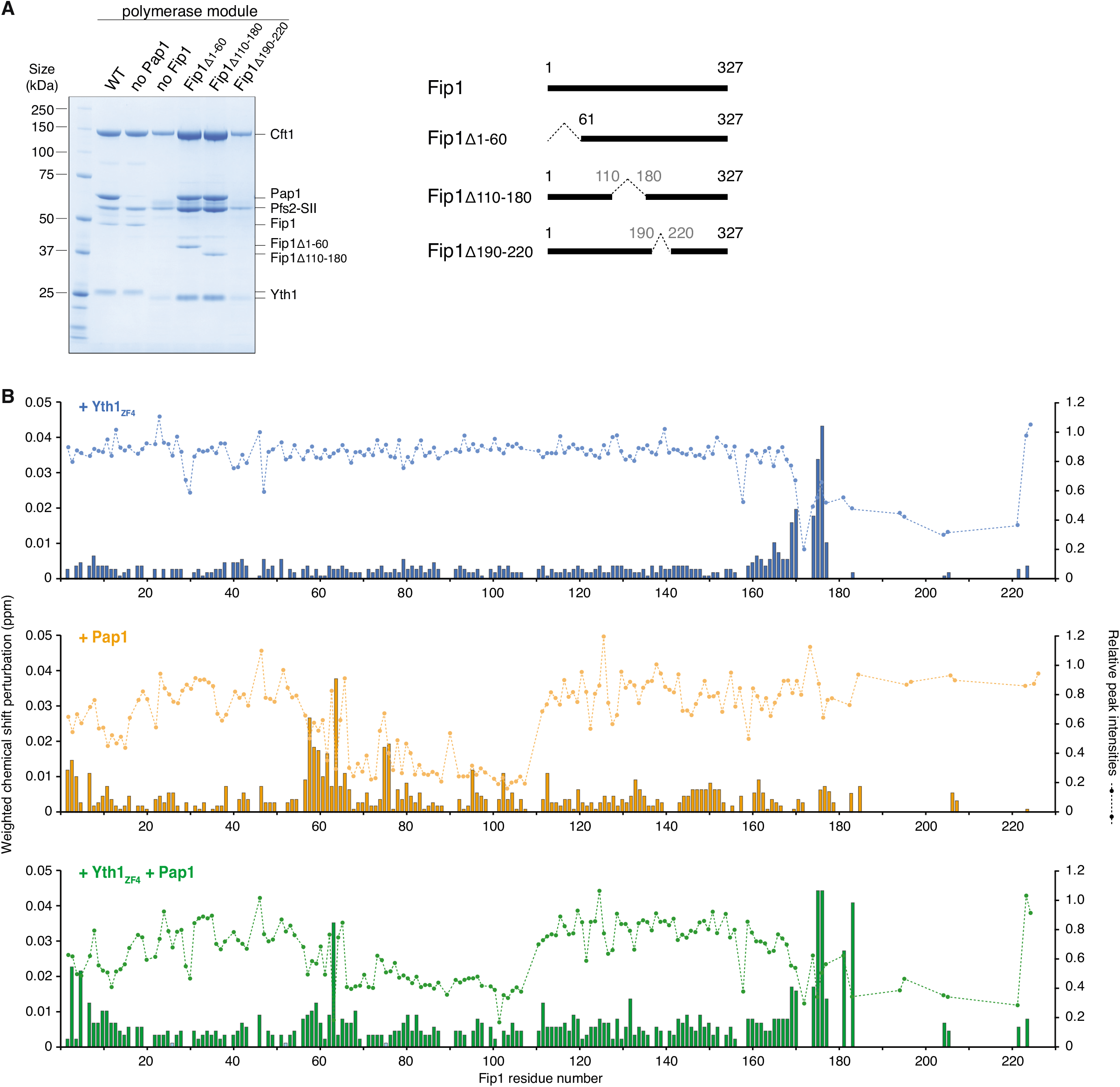
Fip1_226_ contains bipartite binding sites for Yth1 and Pap1. **(A)** SDS-PAGE of pulldown assays using StrepII-tagged (SII) Pfs2. Residues 190–220 on Fip1 are essential for reconstitution of the polymerase module. Yth1 is 6His-tagged for wild-type (WT) and ‘no Pap1’ samples, but untagged in all other samples. Diagrams of full-length and truncated Fip1 are shown on the right. The first three lanes (WT, no Pap1 and no Fip1) are reproduced from Fig. 1. **(B)** Chemical shift perturbations of Fip1_226_ upon binding of Yth1_ZF4_ (blue, top), Pap1 (orange, middle), and Yth1_ZF4_ and Pap1 together (green, bottom). Chemical shift perturbations are shown as histograms. The dotted line indicates the relative peak intensities compared to free Fip1_226_ in the ^1^H,^15^N 2D HSQC spectra. Experiments were performed at 150 mM NaCl to minimize potential non-specific binding.

Next, we used NMR to gain residue-level insight into the molecular interactions of both Yth1 and Pap1 with Fip1 (Fig. 3B, Supplemental Fig. S4A). We incubated Fip1_226_ with Yth1_ZF4_, Pap1, or Yth1_ZF4_ and Pap1 together, and mapped the changes in the spectra. First, upon incubation with Yth1_ZF4_, major chemical shift perturbations and substantial line-broadening were observed for resonances corresponding to Fip1 residues 170–220, indicating a potential interaction between this region and Yth1_ZF4_ (Fig. 3B, top). This region had also exhibited intrinsic line-broadening in the absence of Yth1_ZF4_ (Supplemental Fig. S3D-E). Therefore, we employed ^13^C-detect CON studies using deuterated Fip1_226_. Although the signal intensities were reduced in the ^13^C-detect experiments due to the lower gyromagnetic ratio of ^13^C, line-broadening was reduced by avoiding proton detection. These experiments provide an additional advantage of improved chemical shift dispersion for intrinsically-disordered proteins. We could then unambiguously define additional signals for C-terminal residues (Supplemental Fig. S4B), providing an independent confirmation of our findings. Together, these data revealed that the major Yth1-binding site on Fip1 is within residues 170–220.

Next, upon incubation with unlabeled 66-kDa Pap1, chemical shift perturbations were observed for signals from Fip1_226_ residues 58–80 and signal attenuation as a result of line-broadening was observed for resonances from residues 66–110 (Fig. 3B, middle). In addition, some resonances from the N-terminal acidic region (Fig. 2A) were slightly perturbed upon Pap1 binding, which may be the result of charge-charge interactions. Together, our observations are in agreement with binding of Fip1 residues 80–105 to Pap1, as observed in the crystal structure (Meinke et al. 2008), but our results indicate that additional Fip1 residues (58–110) participate in the interaction. Interestingly, previous studies had shown that Pap1 has higher affinity for full-length Fip1 than for residues 80–105 (Meinke et al. 2008), consistent with involvement of additional residues in this interaction.

Finally, when both Yth1_ZF4_ and Pap1 were added to labelled Fip1_226_, the binding patterns of each individual protein were retained (Fig. 3B, bottom). This suggests that Fip1 contains two independent binding sites – one for Yth1 and one for Pap1. Interestingly, the central LCR was not involved in either of these interactions. Most of the central LCR remains unperturbed, suggesting that this region is still largely disordered and highly dynamic, even when Fip1 is bound to Pap1 and Yth1.

### Reconstitution of a fully recombinant CPF

Residues 110–170 within the central LCR of Fip1 remain dynamic in the presence of Yth1_ZF4_ and Pap1, raising the possibility that they could also be dynamic in the context of the full CPF complex. To investigate this and dissect the structural nature of Fip1 in CPF, we established a strategy to assemble a fully recombinant CPF complex containing all fourteen subunits with selectively-labelled Fip1_226_ that could be used for NMR analysis. Previous biochemical studies used native CPF purified from yeast (Casanal et al. 2017). The recombinant system greatly simplifies genetic manipulation of CPF.

We used a modified version of the biGBac system (Weissmann et al. 2016; Hill et al. 2019) to produce two bacmids: one bacmid contained genes encoding the eight subunits of the nuclease and polymerase modules, and a second bacmid contained the six genes encoding the phosphatase module (Supplemental Fig. S5A). These two bacmids were used for co-infection of *Sf9* cells and the complex was purified using a StrepII-tag on the Ref2 subunit. The isolated complex was then subjected to anion exchange chromatography followed by size exclusion chromatography (Supplemental Fig. S5B). SDS-PAGE analysis of the purified CPF showed the presence of all 14 subunits (Fig. 4A, Supplemental Fig. S5C). Size exclusion chromatography coupled with multi-angle light scattering (SEC-MALS) revealed a molecular weight of 879 ± 28 kDa, which is in agreement with the theoretical molecular weight of CPF (860 kDa) with all subunits in uniform stoichiometry (Supplemental Fig. S5D). We also tested *in vitro* cleavage and polyadenylation activities to determine whether the complex is functionally active. Recombinant CPF specifically cleaves the 3’-UTR of a model pre-mRNA and polyadenylates the cleaved RNA product (Fig. 4B). Thus, we were able to purify a fully recombinant, active CPF.

**Fig. 4.**
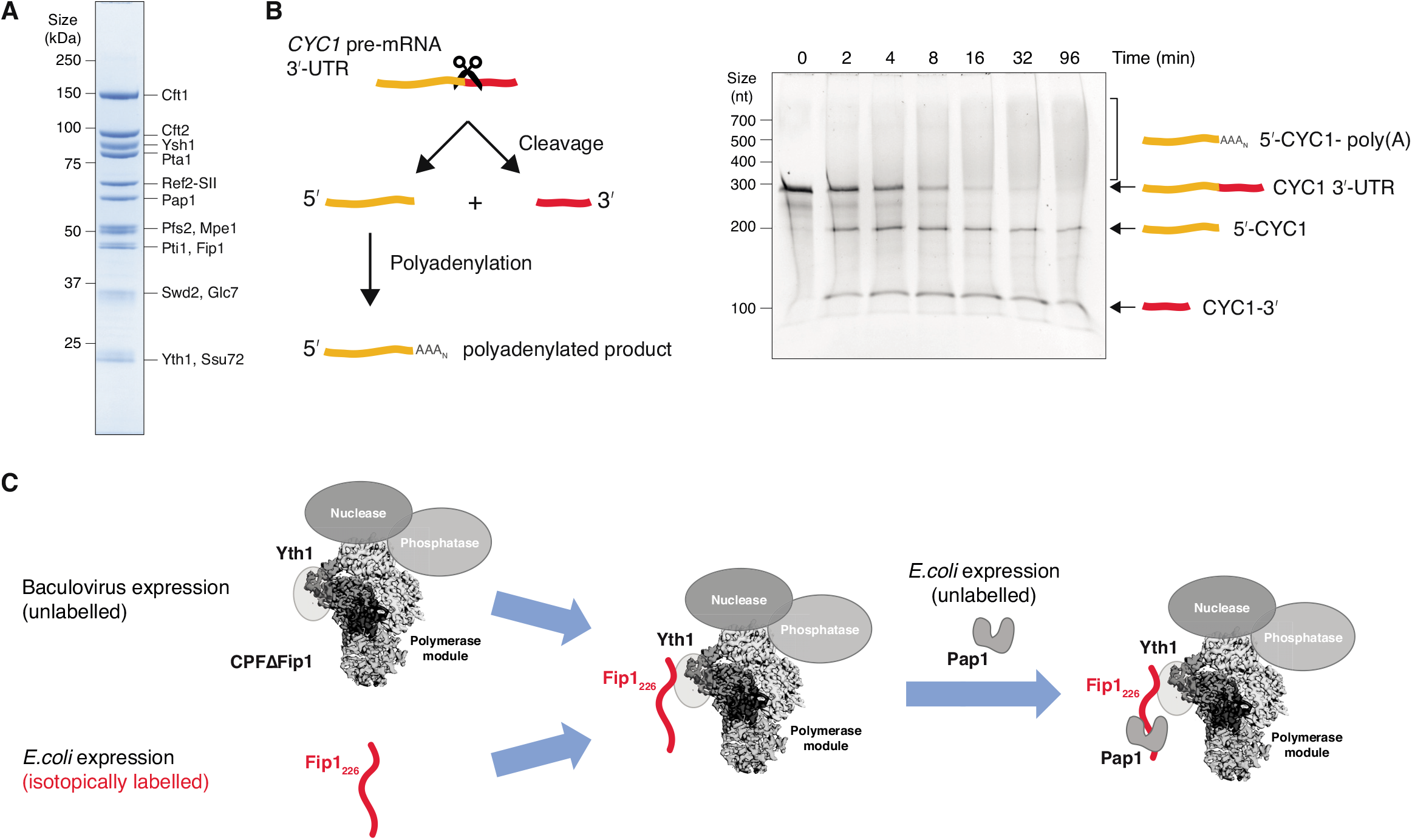
Purification of a recombinant, active CPF. **(A)** SDS-PAGE showing all 14 subunits present in purified recombinant CPF. **(B)** Schematic diagram (left) and denaturing urea polyacrylamide gel (right) of cleavage and polyadenylation assay of recombinant CPF. A 259-nucleotide *CYC1* 3’ UTR was used as a model pre-mRNA substrate. This gel is representative of experiments performed two times. **(C)** Workflow for preparing CPF with selectively-labelled Fip1_226_. Recombinant CPFΔFip1 was purified from baculovirus expression, while isotopically labelled Fip1_226_ was purified after overexpression in *E. coli.* Purified Fip1_226_ was combined with excess CPFΔFip1 and the resulting complex was used for NMR analysis. Free CPFΔFip1 is silent in NMR experiments as it is unlabeled. Excess Pap1 was added to study its interaction with Fip1 on CPF.

Next, we produced a variant of CPF lacking Fip1 (referred to as CPFΔFip1) using the same method (Supplemental Fig. S5E). Notably, Pap1 was also absent, showing that Fip1 is also essential for Pap1 incorporation into intact CPF. We separately expressed and purified isotopically-labelled Fip1_226_ from *E. coli.* To make a selectively-labelled CPF-Fip1_226_ chimeric complex for NMR analysis, we mixed unlabeled CPFΔFip1 in 1.1-fold excess with isotopically-labelled Fip1_226_ (Fig. 4C). The small excess of CPFΔFip1 is not visible by NMR and therefore does not contribute to signals observed in the NMR experiments. Finally, we also added excess Pap1 to study the interaction between Pap1 and CPF-Fip1_226_.

To monitor the integrity and stoichiometry of these complexes, we used mass photometry. By measuring the light scattered by single molecules, mass photometry can be used to determine molecular mass with minimal amounts of protein (10 μl of 100 nM samples) (Young et al. 2018). Mass photometry reported a molecular mass in the range of ~850 kDa for recombinant CPF (Supplemental Fig. S5F), which is in agreement with the expected mass and the SEC-MALS measurement (Supplemental Fig. S5D). Additionally, the reported molecular masses for CPFΔFip1, CPF-Fip1_226_ and CPF-Fip1_226_-Pap1, are all in good agreement with their expected molecular masses (Supplemental Fig. S5F), confirming stable reconstitution of these complexes. These data show that we were able to generate stable NMR samples with ~10 μM selectively-labelled Fip1_226_-CPF.

### Fip1 is dynamic within CPF

To investigate the dynamics of Fip1 when it is incorporated into CPF, we used NMR to study the recombinant complex. As this complex is ~1 MDa in size, we used a combination of ^2^H,^13^C,^15^N selectively-labelled samples and BEST ^1^H,^15^N-TROSY experiments to enhance sensitivity. Spectra of CPF-Fip1_226_ were compared to spectra of free Fip1_226_ at the same concentration of 11 μM (Fig. 5A). Strikingly, even in the spectra of the 850 kDa complex, Fip1_226_ signals can be clearly observed, indicating that Fip1 remains highly dynamic when bound to CPF. To ensure the signals in the CPF-Fip1_226_ spectra corresponded to CPF-bound Fip1_226_ and not to free Fip1_226_, ^15^N-edited ^1^H diffusion experiments were used to determine the diffusion coefficients of the ^15^N-labelled species in the samples (Supplemental Fig. S6). The measured diffusion coefficient of Fip1_226_ alone (4.6 x 10^-11^ m^2^s^-1^) is consistent with a highly mobile, free protein, whereas the diffusion coefficient of CPF-Fip1_226_ (2.8 x 10^-11^ m^2^s^-1^) is consistent with a larger, more slowly diffusing CPF-bound species. This is further supported by pulldowns and mass photometry (Supplemental Fig. S5D-F), which also showed that the CPF-Fip1_226_ complex is stable and intact during NMR data acquisition.

**Fig. 5.**
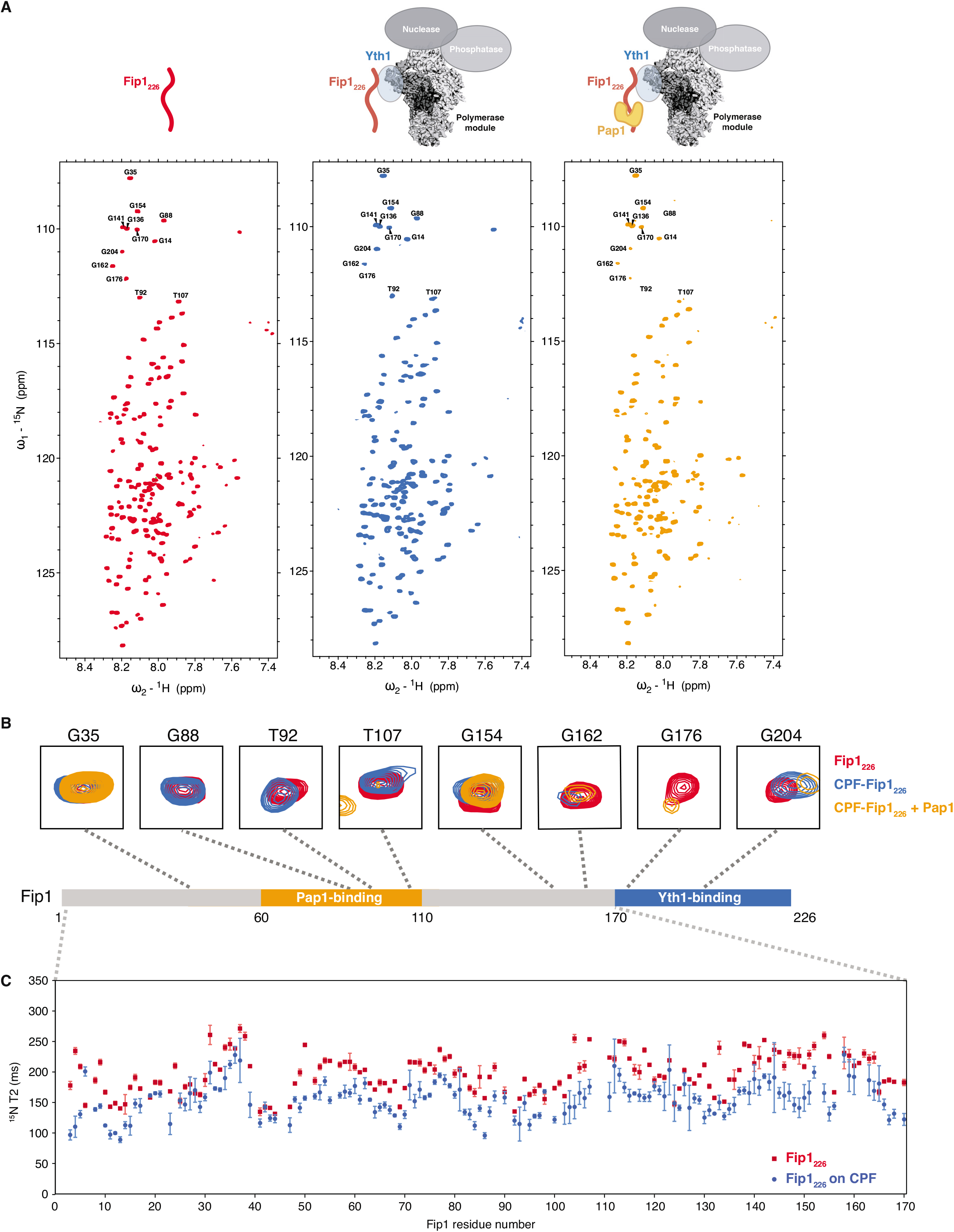
Fip1_226_ is highly dynamic within the CPF complex. **(A)** ^1^H,^15^N 2D HSQC of Fip1_226_, alone (left, red) and bound to CPFΔFip1 (middle, blue) or CPFΔFip1 and Pap1 (right, orange). Schematic diagrams show the proteins included in each experiment. All spectra were collected at 950 MHz with 11 μM ^13^C,^15^N,^2^H Fip1_226_ in 150 mM NaCl buffer. Peaks analyzed in Fig. 5B are indicated in the spectra. **(B)** Selected Fip1_226_ peaks for all three samples. Perturbation or line broadening of peaks specific to the defined regions for Yth1- and Pap1-binding was observed upon interaction with CPFΔFip1 and Pap1. Peaks are mapped onto a diagram of the Fip1_226_ protein. Colors as in (A). **(C)** T2 relaxation data for the first 170 residues of Fip1_226_ alone (red) and incorporated into CPFΔFip1 (blue).

In general, the narrow dispersion of proton chemical shifts in spectra of CPF-Fip1_226_ and free Fip1 226, and the relatively narrow linewidths suggest high local flexibility. This is consistent with Fip1_226_ being largely disordered, both when it is free in solution and when it is incorporated into CPF. In the CPF-Fip1_226_ spectra, we observed selective line-broadening for resonances from residues 170–226, which includes the Yth1-binding region of Fip1 (Supplemental Fig. S7A). For example, the peaks for G176 become much weaker and G204 undergoes chemical shift perturbation when Fip1_226_ is incorporated into CPF (Fig. 5B). This indicates that Fip1_226_ interacts with Yth1 via the interaction site described above to form a stable complex with CPFΔFip1. Other regions of Fip1 are relatively unaffected by incorporation into CPF.

When excess Pap1 was added to CPF-Fip1_226_, selective line-broadening was also observed for resonances in the Pap1-binding region (residues 60–110) (Supplemental Fig. S7A). For example, resonances for G88, T92 and T107 disappear or undergo chemical shift perturbation upon addition of Pap1 (Fig. 5B). This confirms that, similar to the isolated proteins (Fig. 3B), interaction between Pap1 and Fip1 within CPF likely extends beyond that observed in the previously reported crystal structure (Meinke et al. 2008).

Outside the Yth1- and Pap1-binding sites, sharp resonances are present in the CPF-Fip1_226_-Pap1 spectra. These likely represent highly flexible residues in Fip1 and suggest that Fip1_226_ does not have any additional major interactions with CPF. To investigate its dynamics, we determined the ^15^N transverse relaxation times (T_2_) for free Fip1_226_, Fip1_226_-Yth1_ZF4_ and CPF-Fip1_226_ (Fig. 5C). Resonances from the residues in the Yth1-binding region show substantial line-broadening and were therefore not included in this analysis. We found that the interaction of Fip1 with CPF or free Yth1_ZF4_ does not substantially alter the flexibility of the first 170 residues of Fip1_226_ (Fig. 5C, Supplemental Fig. S7B). We also analyzed the flexibility of Fip1_226_ in the presence of Pap1 (Supplemental Fig. S7B). This showed that the Pap1-binding site becomes more rigid upon Pap1 binding, but the residues outside the Pap1-binding site remain highly dynamic. In conclusion, Fip1 is dynamic in the context of CPF.

### The central low-complexity region of Fip1 plays a role in cleavage and polyadenylation

The dynamic central LCR of Fip1 (residues 110–180) between the Pap1- and Yth1-binding sites is of particular interest because it may flexibly tether Pap1 to CPF. This would be consistent with the flexible position of Pap1 in cryoEM analysis of the polymerase module (Fig. 1A). To identify whether the central LCR is functionally important for the cleavage and polyadenylation activities of CPF, we deleted Fip1 residues 110–180 and purified a CPF(Fip1Δ110–180) complex. SDS-PAGE analysis of purified CPF(Fip1Δ110–180) showed that it contained all fourteen subunits in similar stoichiometry to wild-type CPF (Fig. 6A). Thus, the central LCR of Fip1 is not required for assembly of CPF.

**Fig. 6.**
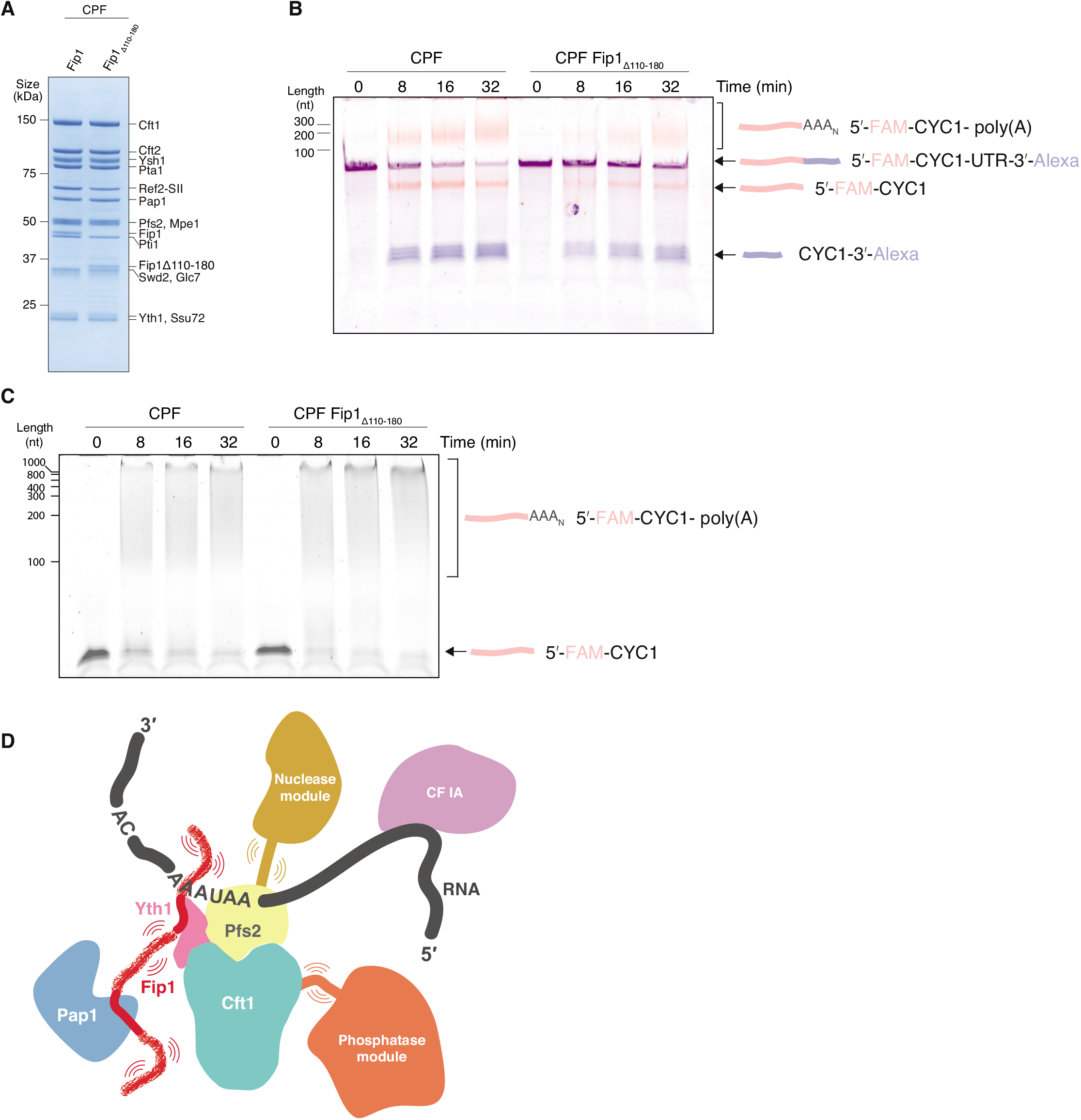
The central LCR of Fip1_226_ is important for cleavage and polyadenylation activities. **(A)** SDS-PAGE of purified CPF alongside CPF(Fip1Δ110-180). Deletion of the central LCR in Fip1 does not affect the assembly and purification of CPF. The composition and stoichiometries of CPF subunits are similar in both samples. **(B)** Denaturing gel electrophoresis of coupled cleavage and polyadenylation assays of a *CYC1* 3’-UTR containing a 5’-FAM and a 3’-A647 label. Each reaction contained 50 nM CPF or CPF(Fip1Δ110-180), 100 nM *CYC1* 3’-UTR and 300 nM CF IA and IB. The cleavage activity of CPF is compromised in the absence of the central LCR in Fip1. This gel is representative of experiments performed two times. **(C)** Polyadenylation activity of wild-type CPF compared to CPF(Fip1Δ110-180). The RNA contains a 5’-FAM label. This gel is representative of experiments performed two times. **(D)** Model for CPF. IDRs within subunits in each of the modules allow flexibility to permit conformational remodeling.

Next, we performed *in vitro* coupled cleavage and polyadenylation assays with both wild-type and mutant complexes. A synthetic 36-nt *CYC1* 3’ UTR RNA was used as a model pre-mRNA substrate. 5’-FAM and 3’-Alexa647 fluorescent labels allowed visualization of the two cleavage products and the polyadenylated RNA. Recombinant wild-type CPF cleaves the substrate RNA efficiently and adds a poly(A) tail to the 5’ cleavage product (Fig 6B, left). In contrast, more substrate RNA remains unprocessed at all time points in the assay for CPF(Fip1Δ110–180) compared to wild-type CPF (Fig. 6B, right). These results could be explained by slower endonucleolytic cleavage of the pre-mRNA or slower activation of the nuclease in CPF lacking Fip1 central LCR.

To assess whether the polyadenylation activity is also defective, we performed an uncoupled polyadenylation assay. Using a 42-nt synthetic *CYC1* RNA ending at the cleavage site (pc*CYC1*), we found that CPF(Fip1Δ110–180) has similar polyadenylation activity to wild-type CPF (Fig. 6C). Interestingly, polyadenylation defects were observed in a previous study where residues 106–190 were replaced with another IDR sequence (Ezeokonkwo et al. 2011). However, this replacement removed some of the highly conserved residues within Fip1 (Supplemental Fig. S3A) and therefore may have disrupted Yth1 positioning within the complex. Thus, our *in vitro* assays show that, surprisingly, the presence of the central LCR in Fip1 is important for efficient pre-mRNA cleavage but not polyadenylation.

## Discussion

mRNA 3’-end processing by CPF is essential for the production of mature mRNA. Here, using *in vitro* reconstitution and structural studies, we gain new insight into the architecture of CPF. We show that the essential Fip1 subunit contains IDRs that are dynamic, even when Fip1 is incorporated into the full CPF complex.

### Fip1 binds Yth1 zinc finger 4 and Pap1

Fip1 interacts directly with zinc finger 4 of Yth1 (Fig. 1A-B) (Helmling et al. 2001; Tacahashi et al. 2003) and is proposed to bind additional 3’-end processing factors such as Pfs2 and Rna14 (Ohnacker et al. 2000). We identified the Yth1- and Pap1-binding sites within Fip1 using NMR but there were no major changes in Fip1_226_ spectra outside these sequences after Fip1 incorporation into intact CPF. Thus, it appears that Fip1 does not interact with other CPF subunits in this context. Fip1 may acquire new interaction partners, e.g. the Rna14 subunit of CF IA, during CPF activation.

Our previous analysis of native yeast CPF showed that up to two Fip1 and two Pap1 molecules can associate with the complex (Casanal et al. 2017), and purified native CPF contains a mixture of 0, 1 or 2 copies of Fip1 (and Pap1). While our work was in preparation, Hamilton and Tong reported a crystal structure of human CPSF30 zinc fingers 4 and 5 bound to 40 residues of hFIP1 (Hamilton and Tong 2020). In their structure, two hFIP1 molecules are bound to one CPSF30, with the same region of each hFIP1 bound to each zinc finger. Notably, the binding affinity of hFIP1 for CPSF30 zinc finger 4 is ~300-fold stronger than for zinc finger 5 (Hamilton and Tong 2020). In our work, yeast Fip1 also shows a preferential binding towards Yth1 zinc finger 4 with some weak interactions with zinc finger 5 (Fig. 1C). A substantial difference in binding affinities for the two zinc fingers could explain why we see only minor changes in the backbone resonances of Yth1 zinc finger 5 upon Fip1 binding. Biophysical characterization by SEC-MALS, mass photometry and NMR diffusion experiments is consistent with one copy of Fip1 in the recombinant CPF. However, we cannot exclude the possibility of a second Fip1 bound to Yth1 at higher Fip1 concentrations, or a mixture of Fip1 and Pap1 stoichiometries in the purified complex (Supplemental Fig. S5F). Like most aspects of 3’-end processing, the interaction and stoichiometry of Fip1 and Yth1 is likely conserved with that of the human proteins.

### Fip1 flexibly tethers Pap1 to CPF and is important for nuclease activation

The poly(A) polymerase Pap1 was previously known to interact with Fip1 (Preker et al. 1995; Meinke et al. 2008). However, some data had suggested that Pap1 may also contact additional CPF subunits (Murthy and Manley 1995; Ezeokonkwo et al. 2011; Casanal et al. 2017) and it was unclear whether this would be required for stable Pap1 incorporation into CPF. Here, we show that Fip1 is essential for Pap1 association with recombinant CPF, but we do not find any evidence for Pap1 contacting other subunits, at least in the apo complex in the absence of RNA and cleavage factors.

A primary functional role of Fip1 may be to flexibly tether Pap1 to CPF, acting as a non-rigid scaffold for the assembly of a fully functional complex. The binding of Yth1 or Pap1 seems to have minimal effect on the dynamics of the rest of Fip1, including the central LCR. Poly(A) tails are synthesized to a length of ~60 As in yeast and 150–200 As in mammals. We hypothesized that a flexible tether would allow Pap1 to follow the growing 3’-end of the poly(A) tail until all adenosines have been added, while allowing CPF to remain bound to the polyadenylation signal in the 3’-UTR. However, deletion of the central LCR in Fip1 did not substantially affect polyadenylation by CPF. Instead, and surprisingly, it compromised pre-mRNA cleavage (Fig. 6). Thus, Fip1 does not act as a ruler for poly(A) tail length.

Activation of the CPF endonuclease must be highly regulated to prevent spurious cleavage. It is likely that upon RNA binding, conformational changes occur to activate the nuclease and allow RNA to access its active site. Our data suggest that the central LCR is required for efficient nuclease activity. One possibility is that a dynamic Fip1 central LCR is required for conformational rearrangements that occur upon pre-mRNA binding and nuclease activation. Alternatively, the Fip1 central LCR may interact with the accessory Cleavage Factors that are necessary for efficient nuclease activity. Another possibility is that the shortened linker in Fip1Δ110–180 results in Fip1 or Pap1 blocking access to the nuclease active site due to steric constraints. In agreement with this, replacing the central LCR with another IDR sequence (instead of deleting it) does not result in cleavage defects (Ezeokonkwo et al. 2011).

### IDRs may mediate conformational transitions in CPF

Unstructured proteins containing IDRs often bind other proteins to form higher order complexes and mediate cellular processes (Wright and Dyson 1999; Dyson and Wright 2005). Fip1 is not the only subunit in CPF that contains IDRs. For example, Mpe1, Ref2, Yth1 and Pfs2 also contain regions that are predicted to be disordered (Nedea et al. 2003; Nedea et al. 2008; Casanal et al. 2017; Hill et al. 2019). The disordered N-terminus of Yth1 binds to Pfs2 (Casanal et al. 2017) but roles for the other IDRs remain unknown.

One possible function for IDRs within CPF is to mediate some of the multiple conformational transitions that allow endonuclease activation, endonuclease inactivation, polyadenylation and transcription termination. Mpe1, a core subunit of the nuclease module, is likely involved in nuclease activation as it binds directly to Ysh1 (Hill et al. 2019). Mpe1 also contains substantial low-complexity regions and is dynamic within a 500 kDa, 8-subunit subcomplex of CPF (Hill et al. 2019). Ref2 is an intrinsically-disordered protein that is important for activation of CPF phosphatase activity and transcription termination (Nedea et al. 2008; Choy et al. 2012; Schreieck et al. 2014). Combined, this would suggest that each module contains at least one IDR-containing protein that allows the dynamics necessary for remodeling of CPF and co-regulation of its enzymatic subunits.

In this work, we established a recombinant CPF system for the first time. This provides us with the potential to generate variants of CPF components to test additional hypotheses regarding the conformational cycle of CPF. Recombinant CPF was essential for studying the dynamics of Fip1 within CPF, and will allow further studies of the dynamics of single proteins within this large, multiprotein complex. Dynamics within large multiprotein complexes, and specifically IDRs, are difficult to study, especially because characterization of mobile regions is often elusive in cryoEM and X-ray crystallography studies. On the other hand, NMR has an advantage in studying dynamic proteins, but for multiprotein complexes, the large molecular masses and overlapping signals from various components pose a major challenge. Our strategy of using a fully recombinant megadalton CPF with selectively-labelled Fip1 ensures the rest of the complex remains NMR-silent, and therefore allows a clean and detailed analysis of a single protein within a large complex. Flexible residues in large complexes can thus produce sharp resonances in NMR spectra, opening up new possibilities to dissect the dynamics and functional roles of IDRs within biological assemblies.

## Materials and Methods

### Bioinformatics analysis

Disorder prediction was performed using IUPRED2A in long disorder form (Meszaros et al. 2018). Net charge per residue and fraction of charged residues were calculated using the localCIDER package with a window size of 10 residues (Holehouse et al. 2017). Sequences of homologs of yeast Fip1 were collected from UniProt database and aligned using ClustalOmega with default parameters (Madeira et al. 2019). The homologs were then divided into N-terminal domain (NTD) and C-terminal domain (CTD) based on the alignment. Residues in the alignment corresponding to yeast Fip1 residues 1–226 were classified as NTD and the rest as CTD. Amino acids were grouped by negatively-charged residues (DE), positively-charged residues (RK), amines (NQ), small hydrophilic residues (ST), aromatic residues (FYW), aliphatic residues (LVIM) and other small residues (PGA). The frequency of occurrence for each group is defined as the total number of residues in a specific group normalized by the length of each sequence. The occurrence of histidines and cystines are minimal and hence omitted from the plot. Visualizations of data were performed either using custom-written python or R scripts.

### DNA constructs

Cloning involving pACEBac1, pBIG1 or pBIG2 was performed in DH5α or TOP10 *E. coli* cells. DH10 EmBacY *E. coli* cells were used to generate and purify all bacmids used in this study. All the CPF subunit genes were synthesized by GeneArt with their sequences optimized for expression in *E. coli.*

The five-subunit wild-type polymerase module was cloned into the MultiBac protein complex production platform as previously described (Casanal et al. 2017). In brief, Cft1, Pfs2-3C-SII and 8His-3C-Yth1 were cloned into the pACEBac1 plasmid. The Pfs2 subunit had a C-terminal 3C protease site and a twin strep tag (SII). The genes for Pap1 and Fip1 were cloned into the pIDC vector. The five-subunit wild-type polymerase module was generated by Cre-Lox recombination as described (Stowell et al. 2016; Casanal et al. 2017).

Polymerase module truncations and deletions were cloned using a modified version of the biGBac system (Weissmann et al. 2016) as described previously (Hill et al. 2019). Yth1_ΔZF45C_, Yth1_ΔZF5C_, Yth1Δ_ZF4_, Fip1_Δ1-60_ and Fip1_226_ were amplified by PCR using primers listed in Supplemental Table S1. For deletion of Yth1 zinc finger 4 and Fip1_Δ110-180_, overlap extension splicing PCR was used. Variants were cloned into pACEBac1 by using BamHI and XbaI restriction sites. The five subunits of the polymerase module were then PCR amplified from their parent plasmid (pACEBac1 for Cft1, Pfs2-3C-SII and Yth1, and pIDC plasmid for Pap1 and Fip1) using biGBac primers as described (Weissmann et al. 2016). Each of the five amplified PCR products therefore contains the individual gene with its own promoter and terminator sequences. The five PCR products were cloned into pBIG1a using Gibson assembly to generate a polymerase module containing a Yth1 or Fip1 variant. The final plasmids were verified by SwaI digestion to ensure that the clone contained all five genes in uniform stoichiometry. For the ΔFip1 construct, the Fip1 PCR product was omitted. For Pap1 purification, the Pap1 gene was cloned into pACEBac1 in-frame with a C-terminal SII tag followed by a 3C protease cleavage site and inserted into pBIG1a.

For the phosphatase module, Pta1 was cloned into pBIG1a, and Ssu72, Pti1, Glc7, Ref2-3C-SII and Swd2, were cloned into pBIG1b by Gibson assembly. The Pta1 gene cassette from pBIG1a and the multi-gene cassette from pBIG1b were released by PmeI digestion. Using a second Gibson assembly step, the PmeI digested gene cassettes were introduced into linearized pBIG2ab. Correct insertion of the six phosphatase module genes into pBIG2ab was verified by SwaI and PacI restriction digestion.

The combined nuclease and phosphatase module (‘CPFcore’) was assembled without any affinity tags in this work. First, the five-subunit polymerase module (Cft1, Pap1, Pfs2, Fip1 and Yth1) was cloned into pBIG1a and a three-subunit nuclease module (Cft2, Ysh1 and Mpe1) was cloned into pBIG1b with Gibson assembly. The multi-gene cassettes from pBIG1a and b were cut by PmeI restriction digestion and cloned into pBIG2ab. For CPFΔFip1, Fip1 was omitted from the CPFcore construct. For Fip1Δ110-180, the Fip1 variant was used.

CF IA subunits Rna14, Rna15, Pcf11 and Clp1 were cloned into the pBIG1c vector using Gibson assembly as described above for the polymerase module subunits.

The sequence corresponding to Fip1 residues 1–226 was cloned into pET28a+ using PCR with primers listed in Supplemental Table S1 and NdeI and HindIII restriction sites. The sequence corresponding to Yth1 zinc finger 4 (residues 108–161) or Yth1 zinc fingers 4, 5 and the rest of the C-terminal region (residues 118–208) was cloned into pGEX6P-2 using PCR with primers listed in Supplemental Table S1 and BamHI and EcoRI restriction sites.

### Baculovirus-mediated protein overexpression

Plasmids encoding the protein or protein complex of interest were transformed into *E. coli* DH10 EmBacY cells. Colonies that had successfully integrated the plasmid into the baculovirus genome were picked using blue/white selection methods. A 5 ml overnight culture of the selected colony was set up in 2 x TY media. For pACEBac1 and pBIG1 vectors, 10 μg/ml Gentamycin was used. For pBIG2 vectors, both 10 μg/ml Gentamycin and 35 μg/ml chloramphenicol were used. Bacmids were purified from these cultures using protocols described earlier (Stowell et al. 2016).

A total of 10 μg of bacmid DNA was transfected into 6 wells of 2×10^6^ adherent *Sf9* cells (at 5×10^5^ cells/ml) using the transfection reagent Fugene HD (Promega). 42–72 h post transfection, the viral supernatant was isolated, diluted two-fold with sterile FBS (Labtech) and filtered through a 0.45 μm sterile filter (Millipore). This primary virus could be stored in the dark at 4 °C for up to 1 year. 0.5 ml of the primary virus was then used to infect 50 ml *Sf9* cells in suspension at ~2×10^6^ cells/ml. The infection was monitored every 24 h by taking note of the cell viability, cell count and YFP fluorescence. 48–72 h post infection, the cells usually undergo growth arrest. During this time, one can observe robust fluorescence indicating high levels of protein expression. The supernatant from this suspension culture was harvested by centrifugation at 2000*g* for 10 min. The resulting supernatant or ‘secondary virus’ was filtered using 0.45 μm pore size sterile filter (Millipore) and was used immediately to infect large-scale expression cultures. For large-scale protein expression, 5 ml secondary virus was used to infect 500 ml suspension *Sf9* cells (at 2×10^6^ cells/ml and viability >90%) grown in 2 l roller bottle flask. All *Sf9* cells were grown in insect-EXPRESS (Lonza) media, incubated in 27°C and at 140 rpm. No additional supplements were provided to the media.

For the production of a recombinant fourteen-subunit CPF, the following modifications were made. 5 ml primary virus was used to infect 100 ml *Sf9* cells (at 2×10^6^ cells/ml) grown in suspension in a 500 ml Erlenmeyer flask. 48–72 h post-infection, when the cell count was ~3×10^6^ cells/ml (> 90% viability) and ~80% of the cells exhibit YFP fluorescence, the supernatant or the secondary virus was harvested. For large-scale over-expression, 5 ml of the phosphatase module secondary virus along with 5 ml of the CPFcore (combined nuclease and polymerase modules) secondary virus were used to infect 500 ml *Sf9* cells (at 2×10^6^ cells/ml) grown in suspension in 2 l roller bottle flask. The cells were harvested either at 48- or 72-h post-infection. The exact time of harvest was decided by performing small-scale protein pull-downs as described in the next section. Upon harvesting, the cell pellets were washed once with pre-chilled PBS, flash frozen in liquid nitrogen and stored at −80°C.

#### Small-scale pulldown assays

Small-scale pulldown assays were used to assess protein expression. 0.5 ml secondary virus was used to infect 50 ml *Sf9* cells (2×10^6^ cells/ml, viability > 90%) in a 200 ml Erlenmeyer flask. For 96 h post-infection, ~10^7^ cells were harvested at 24-h timepoints by centrifugation at 2000*g* for 10 min. Cells were flash-frozen in liquid nitrogen and stored at −20°C. All subsequent steps were performed at 4 °C unless otherwise stated. First, the pellets were lysed in 1 ml pulldown lysis buffer and lysed using vortexing (2 min) with glass beads in a 1.5 ml tube. The lysate was clarified by centrifugation for 30 min in a table top centrifuge and at maximum speed. The supernatant was incubated for 2 h with 20 μl Streptactin resin (GE) that had been washed and pre-equilibrated in pulldown lysis buffer. Protein binding was carried out with mixing for 2 h. Unbound proteins were separated from the resin by centrifugation at 600*g* for 10 min. The resin was then washed twice with 1 ml pulldown wash buffer. The bound proteins were eluted with 20 μl pulldown elution buffer for 5 min. The elution fraction was recovered by centrifugation at 600*g* for 10 min. 12 μl eluted proteins was mixed with 4 μl 4x NuPAGE™ LDS Sample Buffer (Thermofisher) and analyzed by SDS-PAGE (4–12% Bis-Tris gradient gel (Thermofisher) with MOPS running buffer, run at 180 V, for 60 min at room temperature).

### Purification of recombinant CPF complexes

The wild type five-subunit polymerase module was expressed and purified as previously described (Casanal et al. 2017). Buffers for CPF purifications are listed in Supplemental Table S2. All steps were performed at 4 °C unless otherwise stated and the following amounts are given for a prep from 2 l cells. Frozen *Sf9* cells pellets were re-suspended in 120 ml CPF lysis buffer. The cells were lysed by sonication using a 10 mm tip on a VC 750 ultrasonic processor (Sonics). Sonication was performed at 70% amplitude with 5 sec on and 10 sec off. The lysate was clarified by ultracentrifugation at 18,000 rpm in a JA 25.50 rotor for 30 min. The clarified lysate was incubated with 2 ml bed volume StrepTactin Sepharose HP resin (GE) that was pre-equilibrated with CPF lysis buffer. Protein was allowed to bind for 2 h in an end-over-end rotor. The unbound proteins were separated from the resin by centrifugation at 600*g* for 10 min. The resin was then washed in a gravity column with 200 ml CPF wash buffer. Elution was performed at room temperature with ten fractions, each with 3 ml of ice chilled CPF elution buffer incubated for five to ten minutes on the gravity column. The eluted fractions were pooled and loaded on to a 1 ml resource Q anion exchange column (GE) that was equilibrated with CPF wash buffer. CPF was eluted from the resource Q column using a gradient from 0.15–1 M KCl over 100 ml. The eluted fractions were assessed by SDS-PAGE. Such a shallow gradient elution across 100 ml aids in the complete separation of the fourteen-subunit CPF complex from subcomplexes. Next, CPF-containing fractions with stoichiometric subunit amounts were pooled and concentrated in a 50 kDa Amicon centrifugal filter (Sigma) at 4000 rpm in a table top centrifuge. 50 μl concentrated CPF sample was polished further by gel filtration chromatography using a Superose 6 Increase 3.2/300 column (GE) with CPF wash buffer at a flow rate of 0.06 ml/min. The peak fractions from the size exclusion step were analyzed by SDS-PAGE. The fractions were concentrated, flash frozen in liquid nitrogen and stored at −80 °C. For biochemical assays, pure CPF containing fractions were used immediately after the anion exchange purification step.

### Purification of Cleavage factors

CF IB was purified as described previously (Hill et al. 2019). Purification of CF IA was carried out essentially as described for recombinant CPF with its corresponding buffers listed in Supplemental Table 2, and with the following modifications. Pooled eluate fractions from StrepTactin Sepharose HP resin was applied to a 5 ml HiTrap Heparin HP (GE) column equilibrated in CF IA wash buffer, and subsequently eluted using a linear 0.25–1 M NaCl gradient over 100 ml. Following SDS-PAGE analysis and concentration of pooled fractions, CF IA was further purified by gel filtration using a HiLoad 26/60 Superdex 200pg column in CF IA wash buffer. The peak fractions were assessed by SDS-PAGE for sample purity. During concentration of the pooled fractions showing correct complex stoichiometry, care was taken not to over-concentrate the sample (maximum 7 mg/ml). The concentrated purified protein complex was flash frozen in liquid nitrogen and stored at −80 °C until further use.

### Protein expression and purification in E. coli

Yth1 proteins were expressed in BL21 Star with an N-terminal GST-tag. Isotopically-labelled proteins were overexpressed in M9 media (6 g/l Na_2_HPO_4_, 3 g/l KH_2_PO_4_, 0.5 g/l NaCl) supplemented with 1.7 g/l yeast nitrogen base without NH_4_Cl and amino acids (Sigma Y1251). 1 g/l ^15^NH_4_Cl and 4 g/l ^13^C-glucose were supplemented for ^15^N and ^13^C labelling respectively. Expression was induced with 1 mM IPTG at 22°C for 16 h. Harvested cells were lysed by sonication in buffer A supplemented with 2 μg/ml DNase I, 2 μg/ml RNase A and protease inhibitor mixture (Roche). Proteins were bound to GST resin (GE Healthcare) and eluted in buffer A supplemented with 10 mM glutathione (pH-calibrated). Eluted protein was subjected to 3C protease cleavage and loaded onto a Superdex 75 size-exclusion column pre-equilibrated with buffer A. Fractions containing Yth1 were pooled and concentrated using 3,000 MWCO concentrators (Millipore).

Fip1_226_ was expressed in BL21 Star with an N-terminal His-tag. Isotopically-labelled proteins were overexpressed as described for Yth1 constructs. Perdeuterated proteins were overexpressed in cells with step adaptions in media with 10%, 44% and 78% D_2_O, before switching to 100% perdeuterated media supplemented with 1 g/l ^15^NH_4_Cl and 4 g/l ^2^H,^13^C-glucose. Harvested cells were lysed by sonication in buffer B supplemented with 2 μg/ml DNase I, 2 μg/ml RNase A and protease inhibitor mixture (Roche). Proteins were bound to Ni-NTA resin (GE Healthcare) and eluted with buffer B with 250 mM imidazole (pH-calibrated). Eluted protein was subjected to TEV protease cleavage and loaded onto a Superdex 75 size-exclusion column pre-equilibrated with buffer A used for Yth1 constructs. Fractions containing Fip1_226_ were pooled and concentrated using 10,000 MWCO concentrators (Millipore).

Pap1 was expressed in BL21 Star as an N-terminal His-tagged protein. Expression was induced with 1 mM IPTG at 22°C for 16 h. Harvested cells were lysed by sonication in Pap1 lysis buffer. Proteins were bound to Ni-NTA resin (GE Healthcare) and eluted in 50 mM HEPES pH 8.0, 500 mM NaCl, 300 mM imidazole. Elution was exchanged into buffer containing 50 mM HEPES pH 8.0, 100 mM NaCl, 0.5 mM TCEP, loaded onto a HiTrap Heparin column (GE Healthcare) and eluted with Pap1 buffer. Eluted protein was pooled and loaded onto a Superdex 200 size-exclusion column pre-equilibrated with Pap1 buffer. Fractions containing His-Pap1 were pooled and concentrated using 30,000 MWCO concentrators (Millipore).

### CryoEM of polymerase module

UltraAufoil R1.2/1.3 gold supports (Russo and Passmore 2014) were used to make grids of freshly-purified polymerase module containing Pap1. 3 μl purified protein complex was applied onto glow-discharged gold grids in an FEI Vitrobot MKIII chamber maintained at 100% humidity and 4°C followed by 3 s blot (Whatman filter paper) with a blot force of −10 and vitrification in liquid ethane.

Samples were imaged on a FEI Titan Krios operated at 300 keV and equipped with a Falcon-II direct electron detector. A total of 852 micrographs were acquired at a magnification of 47,000x (corresponding to a calibrated pixel size of 1.77 Å) in linear mode. The total electron dose was ~ 35 e– /Å^2^. The frames were aligned and averaged with MotionCorr (Li et al. 2013) and CTF estimation was performed using Gctf embedded in RELION-2 (Kimanius et al. 2016; Zhang 2016). In total, 628 micrographs were selected for further data analysis. ~4,000 particles were manually picked using a mask diameter of 200 Å and a box size of 140 pixels. 2D classes obtained from these manually picked particles were then used as templates for auto-picking in RELION (picking threshold 0.5, minimum inter particle distance 100 Å). A total of 216,375 particles were extracted with a box size of 160 pixels and subjected to of 2D classification. Further 3D classification and refinement led to a map that was highly similar to our previously determined cryoEM map of Cft1-Pfs2-Yth1 subunits with no additional density that could correspond to Pap1.

### Isothermal Titration Calorimetry (ITC)

Samples were prepared in 50 mM HEPES, pH 7.4, 150 mM NaCl. ITC measurements were performed using a MicroCal iTC200 (Malvern) with 300 μM Yth1_ZF4_ in the syringe and 30 μM Fip1_226_ in the cell. The experiments were conducted at 25°C with 14 injections of 2.6 μl preceded by a small 0.5 μl pre-injection that was not included during fitting. For data analysis, appropriate control heats of dilution of Yth1_ZF4_ injected into buffer was subtracted from the raw data and the result was fitted using a single class binding site model in the manufacturer’s PEAQ software to determine the affinity and stoichiometry of binding.

### SEC-MALS

Recombinant CPF was analyzed using a Heleos II 18-angle light scattering instrument (Wyatt Technology) and Optilab rEX online refractive index detector (Wyatt Technology) at room temperature. 100 μl purified recombinant CPF at 1 mg/ml was loaded onto a Superdex 200 10/300 GL increase column (GE Healthcare) pre-equilibrated with 50 mM HEPES pH 7.4, 150 mM NaCl, running at 0.5 ml/min. The molecular mass was determined from the intercept of the Debye plot using the Zimm model as implemented in ASTRA software (Wyatt Technology). Protein concentration was determined from the excess differential refractive index based on a 0.186 refractive index increment for 1 g/ml protein solution.

### Mass photometry / Interferometric Scattering Microscopy

Measurements were performed with the Refeyn One^MP^ iSCAT instrument using coverslips and sample gaskets carefully cleaned with isopropanol. Samples were diluted in 50 mM HEPES pH 7.4, 150 mM NaCl buffer to around 100 nM and 10 μl was loaded into the gasket well. Data were collected for 1 min at 100 Hz and the resultant movies were analyzed using ratiometric averaging of 5 frame bins. Mass was obtained from ratiometric contrast using a standard curve obtained for proteins of known mass measured on the instrument. This technique has been reported to measure molecular mass up to a precision of 1.8 ± 0.5% (Young et al. 2018).

### *In vitro* pulldown assays

Bait proteins and complexes containing a StrepII-tag were diluted to a concentration of 1.5 μM in 50 mM HEPES pH 7.4, 150 mM NaCl. 100 μl of bait protein was mixed with 40 μl bed volume StrepTactin resin (GE Healthcare) and incubated for 1 h at 4°C. Resins were washed with loading buffer three times and eluted with 6 mM desthiobioitin. Elution was analyzed with a 4–12% gradient SDS gel.

### *In vitro* cleavage and polyadenylation assays

Polyadenylation assays were used to test the functional activity of the polymerase module and its variants, and CPF. A 42 nt pre-cleaved *CYC1* (*pcCYC1*) RNA with a 5’ 6-FAM fluorophore (IDT) was used as a substrate for polyadenylation assays as previously described (Casanal et al. 2017).

For couped assays, the 36 nt *CYC1* RNA substrate contained a 5’ 6-FAM fluorophore (IDT) and a 3’ AlexaFluor 647 (IDT) as in (Hill et al. 2019). Reactions contained 100 nM *CYC1* RNA substrate, 50 nM recombinant CPF or its variants, and 300 nM CF IA and CF IB. The reactions were carried out in 10 mM HEPES pH 7.9, 150 mM KOAc, 2 mM Mg(OAc)2, 0.05 mM EDTA, 2% PEG (v/v) with 1 mM DTT and 1 U/μL RiboLock (Thermo), at 30 °C. The reaction products were analyzed by denaturing 20% acrylamide/7 M Urea PAGE to resolve the cleavage products and 10% acrylamide/7 M Urea PAGE to visualize the polyadenylation bands. The gels were pre-heated at 30 W for 30 min prior to loading the samples and running for 10–20 mins at 400 V. The gels were then scanned on a Typhoon FLA-7000 (GE) using the 473 nm laser/Y520 filter for FAM and the 635 nm laser/R670 filter for AlexaFluor647.

### NMR Spectroscopy

Most experiments on Yth1 and Fip1_226_ were performed using in-house Bruker 700 MHz Avance II+ and 800 MHz Avance III spectrometers, both equipped with triple resonance TCI CryoProbe. For some samples (as indicated below), we also utilized the Bruker 950 MHz Avance III spectrometer located at MRC Biomedical NMR Centre.

All experimental data on Yth1 constructs were collected at 700 MHz in 50 mM HEPES pH 7.4, 150 mM NaCl. ^15^N-labelled proteins were used for binding studies and ^13^C, ^15^N-labelled proteins were used for backbone assignment. Backbone experiments and relaxation experiments were acquired at 278 K to extend sample lifetimes and binding experiments were acquired at 298 K to overcome exchange broadening. The dependency of individual peaks was studied by increasing the temperature in 5 K steps.

Experimental data on Fip1_226_ were collected at 700, 800 and 950 MHz. All experiments on Fip1_226_ were carried out at 278K. Backbone experiments were acquired using ^2^H, ^13^C, ^15^N-labelled samples at 800 MHz and 950 MHz in 50 mM HEPES pH 7.4, 50 mM NaCl to recover most signals from exchange broadening. ^13^C-detect experiments were acquired at 700 MHz. Binding studies, unless otherwise specified, were carried out at 800 MHz in 50 mM HEPES pH 7.4, 150 mM NaCl.

To prepare CPF for NMR, isotopically labelled Fip1_226_ was mixed with 1.1-fold molar excess of CPFΔFip1. The complex was buffer-exchanged into 50 mM HEPES pH 7.4, 150 mM NaCl. His-Pap1 used for binding studies was exchanged into the same buffer before being added to the CPF-Fip1_226_ samples. Experimental data on CPF-Fip226 were collected at 950 MHz. All experiments were carried out at 278K in 50 mM HEPES pH 7.4, 150 mM NaCl. 5% D_2_O and 0.05% sodium azide were added to the samples before NMR analysis.

#### Backbone assignment

Assignment of backbone amide peaks of Yth1 constructs was carried out using the following standard triple resonance spectra: HNCO, HN(CA)CO, HNCA, HNCACB, HN(CO)CACB, HN(CAN)NH and HN(COCA)NNH (Bruker). TROSY versions of these spectra were used for the backbone assignment of Fip1_226_. Backbone datasets were collected with non-uniform sampling at 20–50% and processed with compressed sensing using MddNMR package (Jaravine et al. 2008). Resonances from proline residues in Fip1_226_ were assigned using ^1^H start version of ^13^C-detect CON, H(CA)CON and H(CA)NCO (Bruker). Backbone resonances were assigned manually with the aid of Mars (Jung and Zweckstetter 2004). Topspin 3.6 (Bruker) was used for processing and NMRFAM-Sparky 1.47 (Lee et al. 2015) for spectra analysis.

#### Secondary chemical shifts

Cα/Cβ chemical shift deviations were calculated using the following equation:

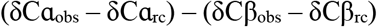

where δCα_obs_ and δCβ_obs_ are the observed Cα and Cβ chemical shifts and δCα_rc_ and δCβ_rc_ are the Cα and Cβ chemical shifts for random coils (Kjaergaard and Poulsen 2011). Temperature coefficients (Kjaergaard et al. 2011) and correction factors for perdeuteration (Maltsev et al. 2012) were applied to the random coil chemical shifts.

#### Binding studies

Weighted chemical shift perturbations were calculated using the equation (Ayed et al. 2001):

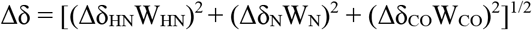

with weight factors determined from the average variances of chemical shifts in the BioMagResBank chemical shift database (Mulder et al. 1999), where W_HN_ = 1, W_N_ = 0.16 and W_CO_ = 0.34.

#### Relaxation measurements

T2 relaxation times were measured using standard INEPT based 3D pulse sequences (Bruker) at a spin lock field of 500 Hz and initial delay of 5 s. 12 mixing times were collected (8.48, 16.96, 33.92, 50.88, 67.84, 101.76, 135.68, 169.6, 203.52, 237.44, 271.36, 8.48 ms) and peak height analysis was done in NMRFAM-Sparky 1.47 (Lee et al. 2015). ^15^N{^1^H}-hetNOE measurements were carried out using standard Bruker pulse programs, with interleaved on- (I) or off-resonance (I0) saturation. The hetNOE values were analyzed in NMRFAM-Sparky 1.47 taking I/I_0_.

#### Diffusion experiments

An ^15^N-edited ^1^H XSTE diffusion experiment with watergate (Ferrage et al. 2003) was used to measure diffusion coefficients of ^15^N-labelled species in the sample, using a diffusion delay of 100 ms and a 4 ms gradient pulse pair for encoding and decoding respectively. Peak intensities at two gradient strengths (5 and 95%) were integrated and the diffusion coefficient was calculated using Stejskal-Tanner equation, where *I* is peak intensity, *G* is gradient strength, *δ* is length of gradient pulse pair, γ is ^1^H gyromagnetic ratio and Δ is diffusion delays:

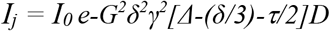

Hydrodynamic radius was deduced using the Stokes-Einstein equation *R_h_ = kT / (6πηD)* where *k* is the Boltzmann constant, *T* is absolute temperature, and *η* is solvent viscosity. The hydrodynamic radius was converted to the effective molecular mass using the equation *R_h_ = 0.066M^1/3^* (Erickson 2009).

## Data Availability

NMR datasets have been deposed in BMRB with accession codes 50795 (Fip1_226_), 50796 (Yth1_ZF4_) and 50797 (Yth1_ZF45C_).

## Acknowledgements

We thank Chris Johnson (MRC LMB) for help with biophysics, Chris Hill (MRC LMB) for CF IA plasmids, and members of the Passmore lab for assistance and advice. This work was supported by a Gates Cambridge PhD Studentship (to A.K.); the European Union’s Horizon 2020 research and innovation programme (ERC grant No. 725685, to L.A.P.); the Marie Skłodowska-Curie grant agreement No. 838945 to X.L.), and the Medical Research Council, as part of United Kingdom Research and Innovation (MRC grant No. MC_U105192715, to L.A.P.). Some of the NMR studies were supported by the Francis Crick Institute through access to the MRC Biomedical NMR Centre. The Francis Crick Institute receives its core funding from Cancer Research UK (FC001029), the UK Medical Research Council (FC001029), and the Wellcome Trust (FC001029).

## Author Contributions

A.K., C.W.H.Y. and L.A.P. conceived the study. A.K., C.W.H.Y. and J.B.R.M. purified proteins and performed assays. C.W.H.Y. performed biophysical assays. C.W.H.Y. and S.M.V.F. performed NMR studies. X.L. performed bioinformatic analyses. A.K., C.W.H.Y. and L.A.P. wrote the manuscript. All authors discussed and commented on the final manuscript.

