## Supplementary Information for "Dynamics in Fip1 regulate eukaryotic mRNA 3'-end processing"

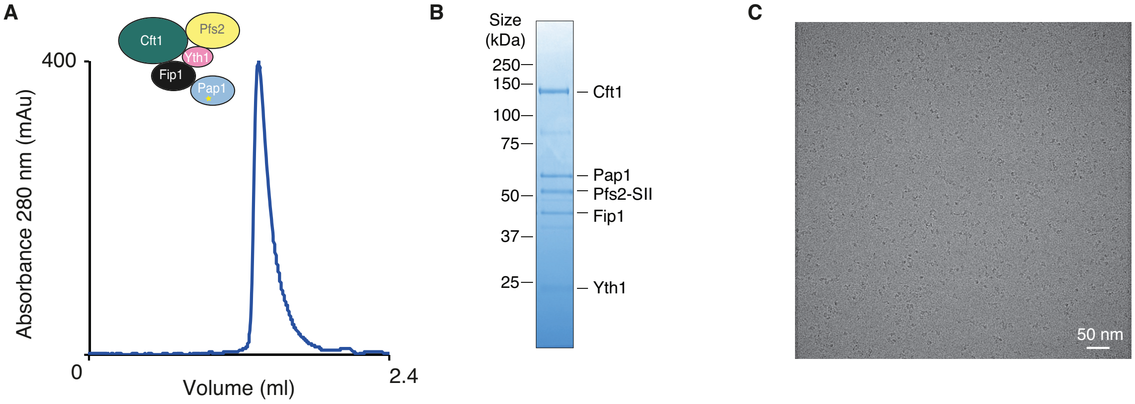


**Supplemental Fig. S1. CryoEM of yeast polymerase module. (A)** Size exclusion chromatogram of purified polymerase module. **(B)** SDS-PAGE showing the five subunits of purified polymerase module. Pfs2 has a StrepII tag (SII). **(C)** Cryo-EM micrograph of five-subunit polymerase module.

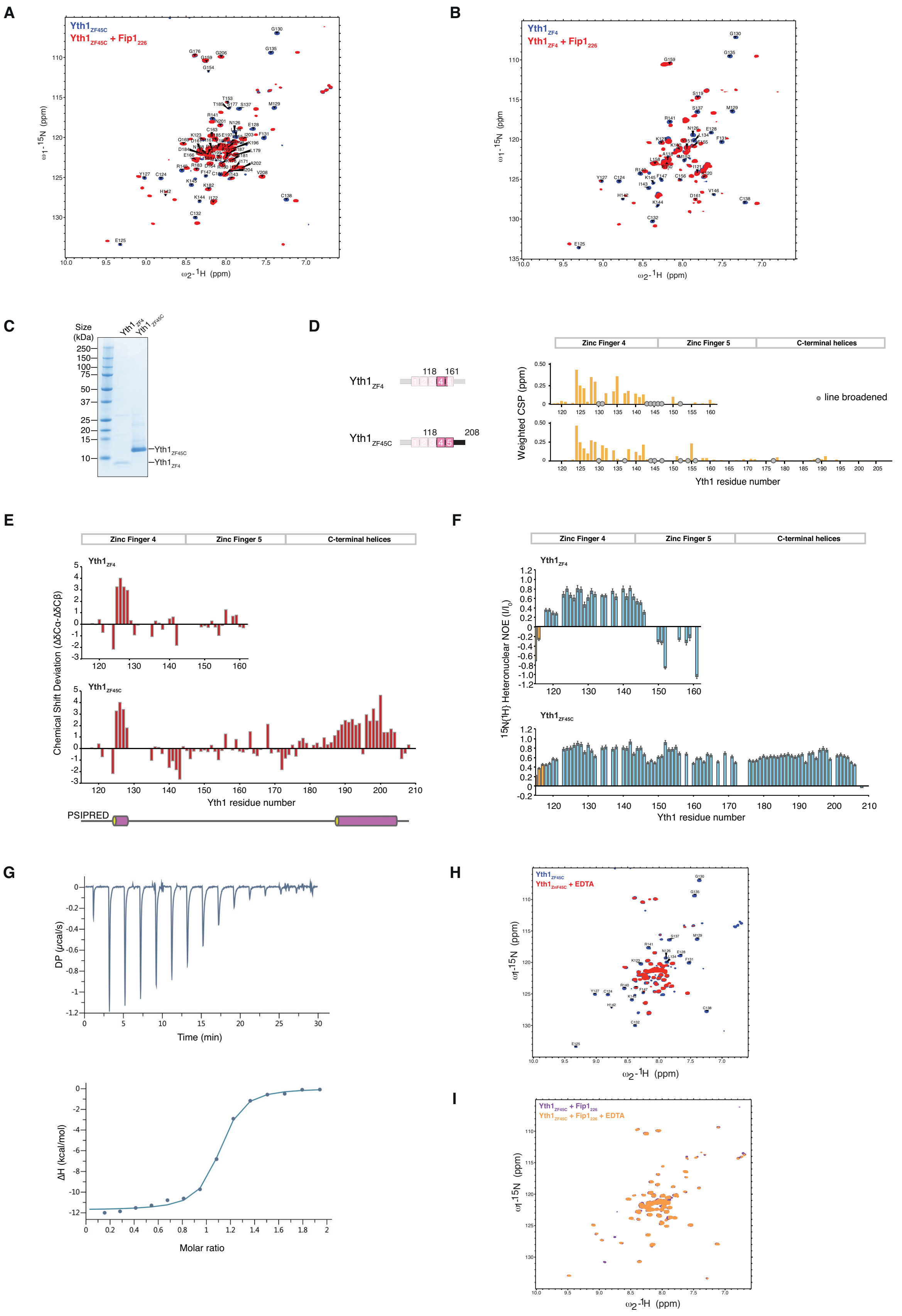

A
Yth1ZF45C
Yth1ZF45C + Fip1226
ω1-15N (ppm)
ω2-1H (ppm)
B
Yth1ZF4
Yth1ZF4 + Fip1226
ω1-15N (ppm)
ω2-1H (ppm)
C
Size (kDa)
250
150
100
75
50
37
25
20
15
10
Yth1ZF4
Yth1ZF45C
D
Yth1ZF4
118
161
Yth1ZF45C
118
208
Zinc Finger 4
Zinc Finger 5
C-terminal helices
Weighted CSP (ppm)
line broadened
Yth1 residue number
E
Zinc Finger 4
Zinc Finger 5
C-terminal helices
Yth1ZF4
Yth1ZF45C
Chemical Shift Deviation (ΔδCα-ΔδCβ)
Yth1 residue number
PSIPRED
F
Zinc Finger 4
Zinc Finger 5
C-terminal helices
Yth1ZF4
Yth1ZF45C
15N{1H} Heteronuclear NOE (I/I0)
Yth1 residue number
G
DP (μcal/s)
Time (min)
ΔH (kcal/mol)
Molar ratio
H
Yth1ZF45C
Yth1ZnF45C + EDTA
ω1-15N (ppm)
ω2-1H (ppm)
I
Yth1ZF45C + Fip1226
Yth1ZF45C + Fip1226 + EDTA
ω1-15N (ppm)
ω2-1H (ppm)

**Supplemental Fig. S2. Fip1 interacts with zinc finger domain 4 of Yth1.** $^{1}H,^{15}N$ 2D HSQCs of **(A)** $Yth1_{ZF45C}$ and **(B)** $Yth1_{ZF4}$ reveal major chemical shift perturbations upon binding to an equimolar amount of $Fip1_{226}$. Backbone assignment of the two Yth1 constructs are labelled on the spectra. **(C)** SDS-PAGE showing the two Yth1 constructs used for NMR analysis. **(D)** Chemical shift perturbations in the HSQC spectra mapped onto the Yth1 sequence. A similar pattern was observed in both constructs, and most of the perturbations are localized in zinc finger 4. Grey circles indicate resonances that showed exchange broadening upon Fip1 binding. **(E)** Cα/Cβ chemical shift deviations reveal secondary structure propensities. Positive values suggest helical propensity and negative values suggest propensity for β-strand formation. PSIPRED prediction for $Yth1_{ZF45C}$ is shown in the schematic below (tube for alpha helix). **(F)** $^{15}N\{^{1}H\}$ heteronuclear NOE values were recorded at 278K and plotted as $I/I_0$. Extra residues at the N-termini of the Yth1 constructs that result from the cloning strategy are colored in orange. Comparison between $Yth1_{ZF4}$ and $Yth1_{ZF45C}$ shows zinc finger 4 retains the properties as in $Yth1_{ZF45C}$ with a possible domain boundary at residue 148. Error bars represent experimental deviations between two hetNOEs experiments. **(G)** Binding affinity of Yth1 and Fip1 was determined by isothermal calorimetry (ITC). 300 μM $Yth1_{ZF4}$ was injected into 30 μM $Fip1_{226}$ and the measured $K_D$ was 240 ± 40 nM. The number of binding sites on $Fip1_{226}$ for $Yth1_{ZF4}$ was determined to be 1.05 ± 0.01 in these conditions. We could not obtain $Fip1_{226}$ in high enough concentration to do the reverse titration and measure the number of Fip1 binding sites on Yth1. **(H)** $^{1}H,^{15}N$ 2D HSQC of $Yth1_{ZF45C}$ with (red) and without (blue) 1 mM EDTA. Crosspeaks from residues 118–142 of $Yth1_{ZF4}$ are labelled in the spectra. EDTA addition results in a spectrum with narrow dispersion in proton chemical shift, suggesting the protein has lost its globular fold. **(I)** $^{1}H,^{15}N$ 2D HSQC of $Yth1_{ZF45C}$-$Fip1_{226}$ with (orange) and without (purple) 1 mM EDTA. The complex was prepared by mixing equimolar amounts of $Yth1_{ZF45C}$ and $Fip1_{226}$, and it remains stable in 1 mM EDTA.

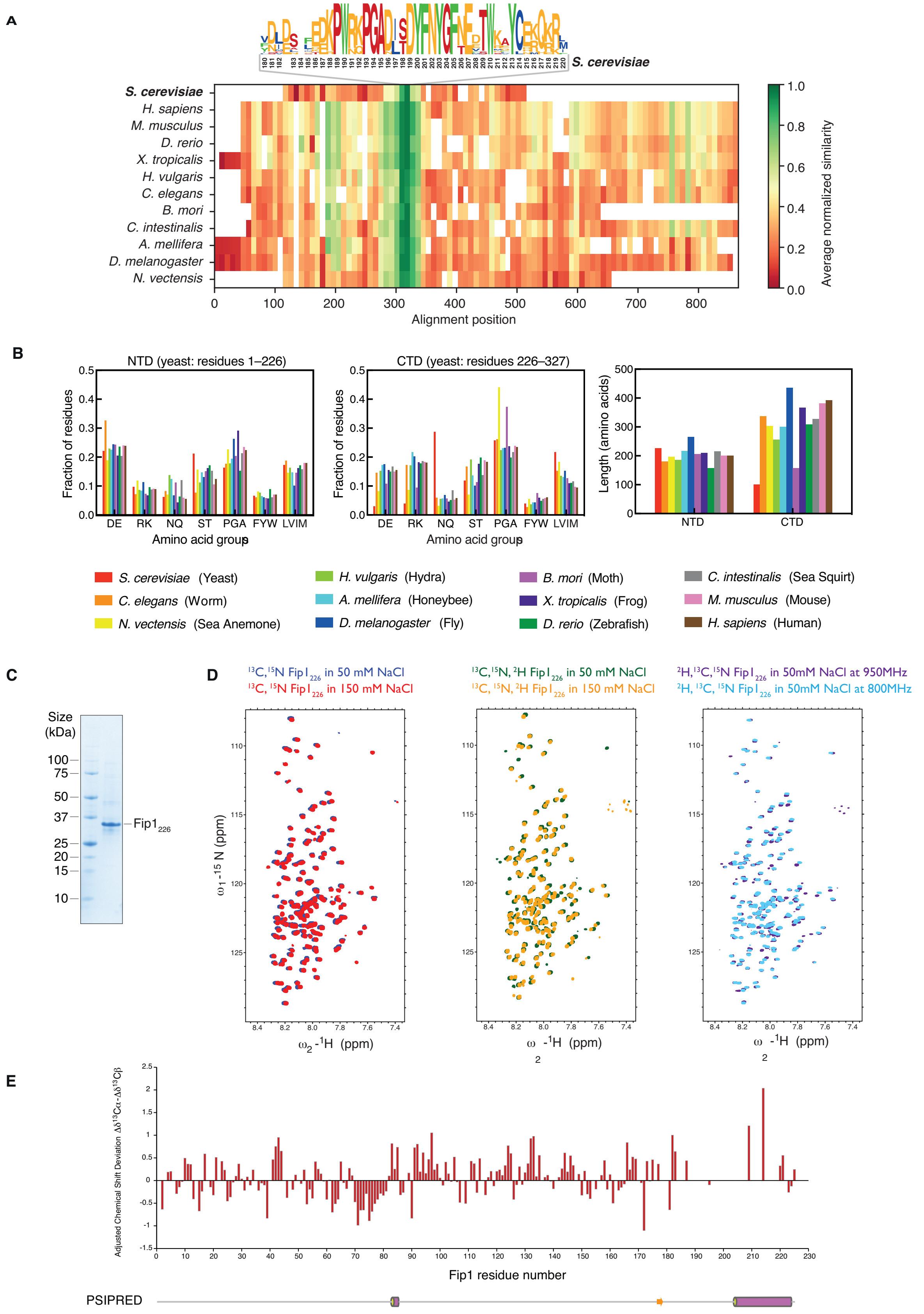

A
S. cerevisiae
H. sapiens
M. musculus
D. rerio
X. tropicalis
H. vulgaris
C. elegans
B. mori
C. intestinalis
A. mellifera
D. melanogaster
N. vectensis
Alignment position
Average normalized similarity
B
NTD (yeast: residues 1–226)
CTD (yeast: residues 226–327)
Fraction of residues
Amino acid group
DE RK NQ ST PGA FYW LVIM
Length (amino acids)
NTD
CTD
S. cerevisiae (Yeast)
C. elegans (Worm)
N. vectensis (Sea Anemone)
H. vulgaris (Hydra)
A. mellifera (Honeybee)
D. melanogaster (Fly)
B. mori (Moth)
X. tropicalis (Frog)
D. rerio (Zebrafish)
C. intestinalis (Sea Squirt)
M. musculus (Mouse)
H. sapiens (Human)
C
Size (kDa)
100
75
50
37
25
20
15
10
Fip1226
D
13C,15N Fip1226 in 50 mM NaCl
13C, 15N Fip1226 in 150 mM NaCl
13C,15N, 2H Fip1226 in 50 mM NaCl
13C, 15N, 2H Fip1226 in 150 mM NaCl
2H,13C,15N Fip1226 in 50mM NaCl at 950MHz
2H, 13C, 15N Fip1226 in 50mM NaCl at 800MHz
ω1 -15 N (ppm)
ω2 -1H (ppm)
E
Adjusted Chemical Shift Deviation Δδ13Cα-Δδ13Cβ
Fip1 residue number
PSIPRED

**Supplemental Fig. S3. Sequence and structure analysis of Fip1. (A)** Plot of sequence homology of Fip1 across various eukaryotic species. High conservation is only observed between residues 190–210, as highlighted in the sequence logo. **(B)** Comparison of amino acid compositions and lengths of the N-terminal domains (NTD, residues 1–226 in yeast) and C-terminal domains (CTD, residues 227–327 in yeast) of Fip1 across species. **(C)** SDS-PAGE showing the purified $Fip1_{226}$ construct used for NMR analysis. **(D)** To retrieve some of the line-broadened signals, $Fip1_{226}$ spectra were compared after collection in different conditions including 50 mM and 150 mM NaCl, with and without $^{2}H$-labelling, and at 800 MHz and 950 MHz. Assignment of backbone resonances was performed using data collected at 800 and 950 MHz with $^{13}C$,$^{15}N$,$^{2}H$ $Fip1_{226}$ in 50 mM NaCl (see Methods). Line-broadening is particularly prevalent for peaks from residues 170–220. **(E)** Plot of Cα/Cβ chemical shift deviations. Positive values suggest helical propensity and negative values suggest β-strand propensity. PSIPRED prediction of secondary structure for $Fip1_{226}$ is shown in cartoon, below the plot.

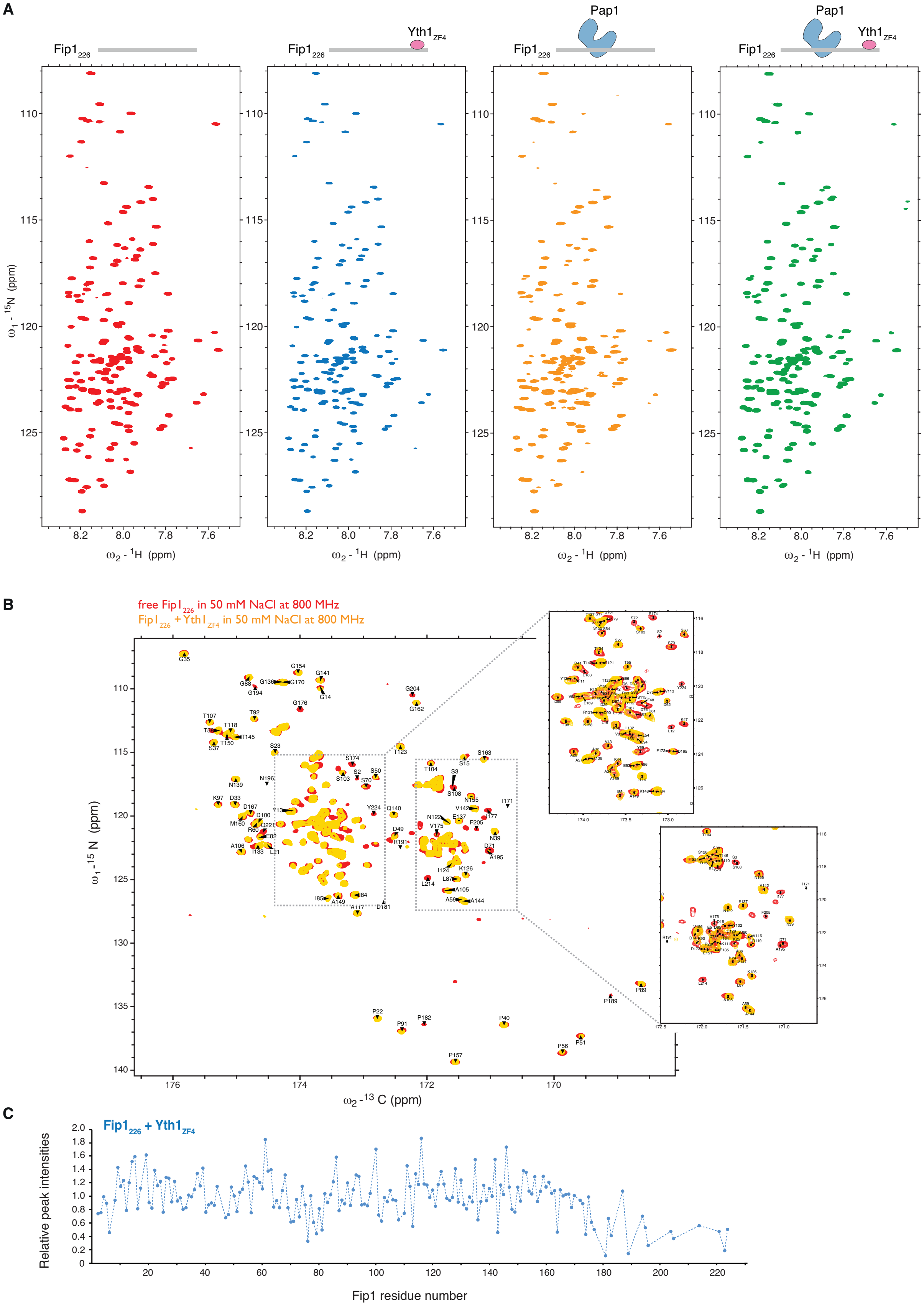

A
Fip1226
Fip1226
Yth1ZF4
Pap1
Fip1226
Pap1
Fip1226
Yth1ZF4
ω1 - 15N (ppm)
ω2 - 1H (ppm)
B
free Fip1226 in 50 mM NaCl at 800 MHz
Fip1226 + Yth1ZF4 in 50 mM NaCl at 800 MHz
ω1 -15 N (ppm)
ω2 -13 C (ppm)
C
Fip1226 + Yth1ZF4
Relative peak intensities
Fip1 residue number

**Supplemental Fig. S4. Fip1$_{226}$ binding studies. (A)** $^{1}H,^{15}N$ 2D HSQC for free Fip1$_{226}$ (red), and Fip1$_{226}$ in the presence of Yth1$_{ZF4}$ (blue), Pap1 (orange), and both Yth1$_{ZF4}$ and Pap1 (green). Cartoons depicting the proteins included in each experiment are shown above. Spectra were collected at 800 MHz with $^{13}C,^{15}N$ Fip1$_{226}$ in 150 mM NaCl buffer. **(B)** $^{13}C$-detect $^{13}C,^{15}N$ 2D CON spectra for free Fip1$_{226}$ (red) and Fip1$_{226}$ in the presence of Yth1$_{ZF4}$ (yellow). To retrieve some of the missing peaks due to exchange broadening, CON spectra were collected at 800 MHz with $^{13}C,^{15}N,^{2}H$ Fip1$_{226}$ in 50 mM NaCl buffer. **(C)** Changes in relative peak intensities for Fip1$_{226}$ in the presence of Yth1$_{ZF4}$, compared to free Fip1$_{226}$.

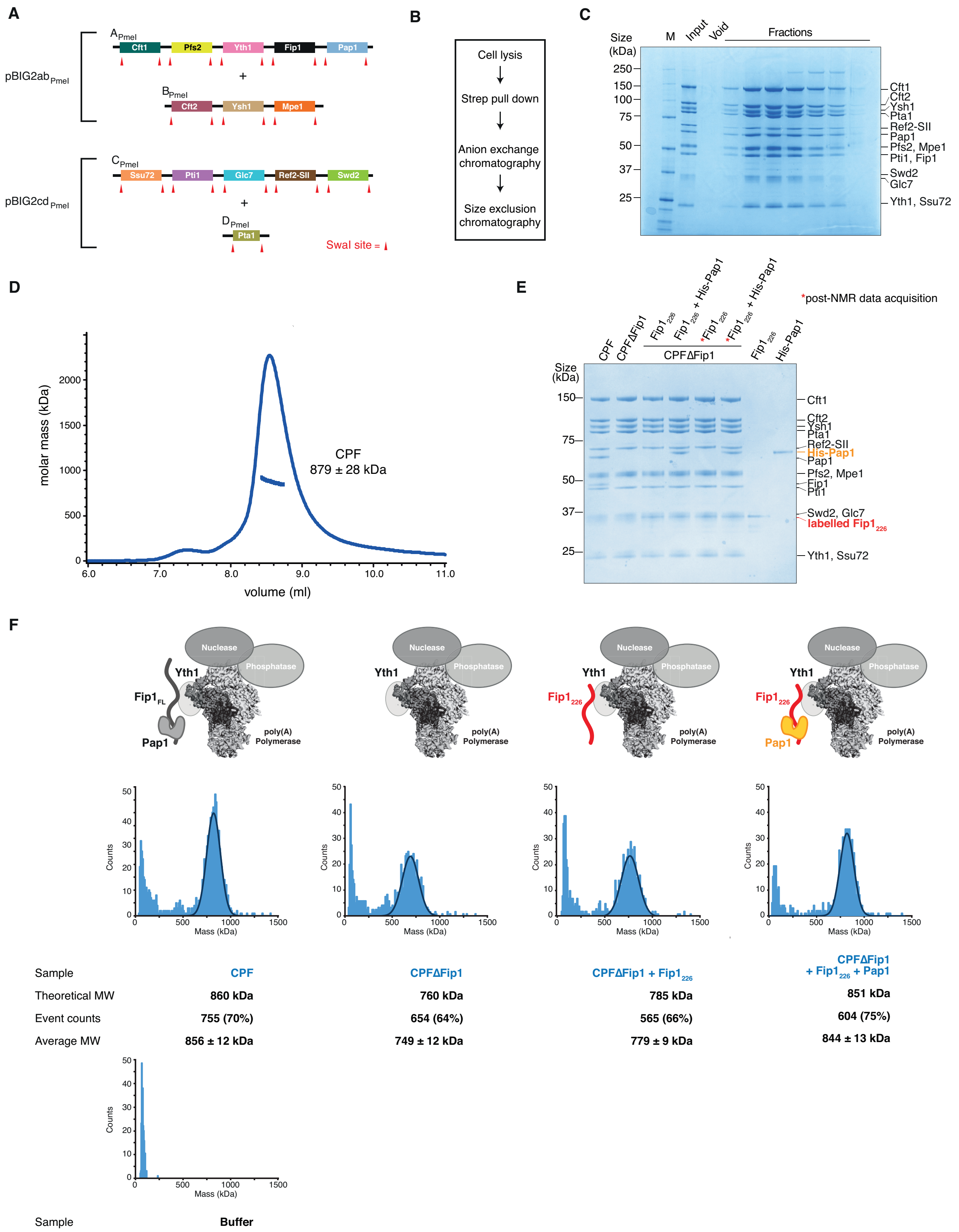


**Supplemental Fig. S5. Purification of recombinant CPF and reconstitution of CPF-Fip1$_{226}$ for NMR analysis. (A)** Multi-bacmid strategy for producing recombinant CPF using a baculovirus expression system. Cartoons of PmeI-digested fragments are shown. The pBIG2ab bacmid contains subunits of the nuclease and polymerase modules, and the pBIG2cd bacmid contains subunits of the phosphatase module. The two bacmids were used for coinfection of *Sf*9 insect cells. **(B)** Workflow for recombinant CPF purification. **(C)** SDS-PAGE showing fractions from size exclusion chromatography of recombinant CPF. M, molecular weight marker. **(D)** SEC-MALS analysis of recombinant CPF. The dominant species has an average molecular weight of 879 ± 28 kDa, which is consistent with the expected molecular weight of 860 kDa. (**E)** Pulldown assays of reconstituted CPF complexes using StrepII-tagged (SII) Pfs2. Both Fip1 and Pap1 are absent in the CPFΔFip1 sample. *E. coli* expressed His-Pap1 and $Fip1_{226}$ are shown on the right. The same samples were analyzed by pull-downs and visualized by SDS-PAGE after NMR data acquisition to confirm sample stability (red asterisks). **(F)** Recombinant CPF complexes were analyzed by mass photometry. Schematic diagrams indicate the sample used in each analysis. The masses obtained for the events observed are plotted as histograms and the peaks are fitted to simple gaussian distributions indicated by the black lines. The number of events within these peaks and the percentage of the total events detected are indicated. Measurements were done in triplicate and the reported error values are the calculated standard deviations. Given the gaussian distributions of masses, we cannot exclude that CPF contains a mixture of complexes with 0, 1 and 2 copies of Fip1 and Pap1. A buffer only blank indicates that it contributes the majority of background events detected with masses below 75 kDa in the CPF complex measurements.

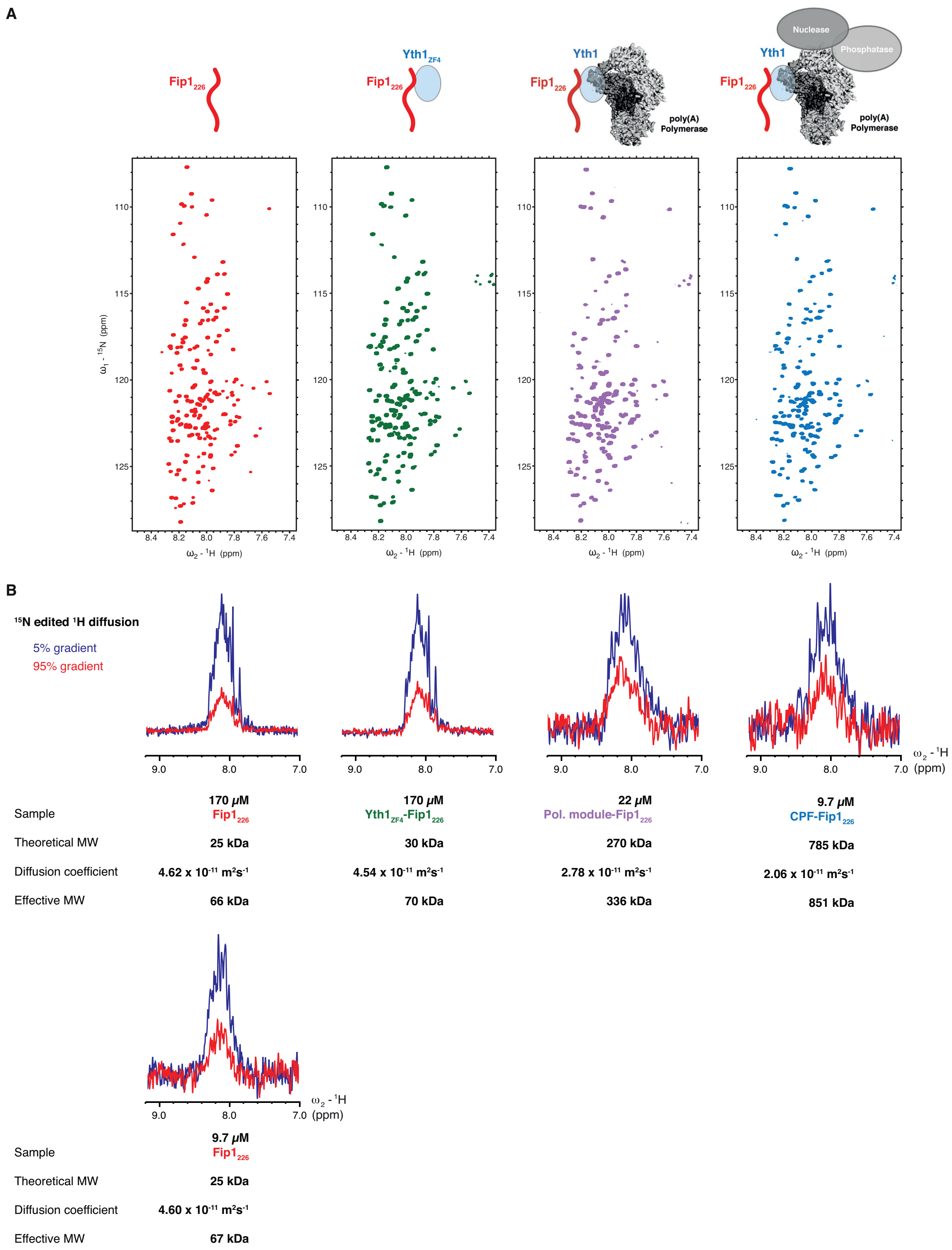


**Supplemental Fig. S6. Diffusion experiments to monitor sample stability and size during NMR data acquisition. (A)** $^{1}$H,$^{15}$N 2D HSQC of free Fip1$_{226}$ (red), Yth1$_{ZF4}$-Fip1$_{226}$ (green), polymerase module-Fip1$_{226}$ (purple) and CPF-Fip1$_{226}$ (blue). Schematic diagrams indicate the protein components in each sample. **(B)** $^{15}$N-edited $^{1}$H diffusion experiments were used to estimate the diffusion coefficients of the $^{15}$N-labelled species in each sample. Smaller proteins diffuse more quickly and have a larger signal attenuation. By measuring the signal attenuation in two gradient fields, the diffusion coefficients were determined and converted to their effective molecular masses. Measurements at two different concentrations of $Fip1_{226}$ (170 and 9.7 μM) show that determination of the diffusion coefficient is reproducible and that the method is sensitive even at low concentrations.

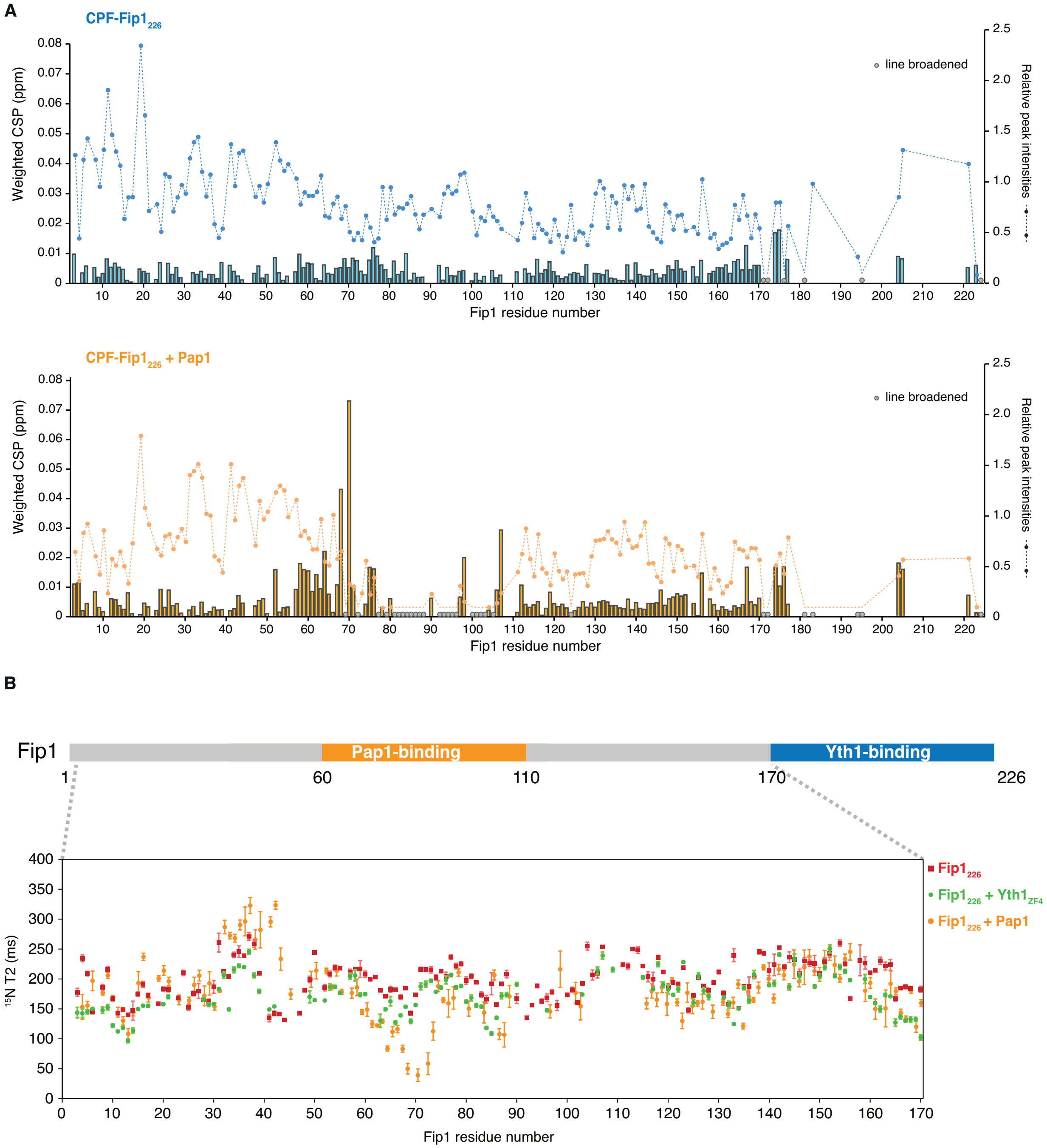


**Supplemental Fig. S7. Chemical shift perturbations and exchange-broadening in CPF-Fip1$_{226}$ binding studies. (A)** Detailed analysis of the HSQC spectra shown in Fig. 5A. Chemical shift perturbations (CSPs) are shown as histograms. Relative peak intensities compared to free Fip1$_{226}$ at the same concentration (11 µM) are indicated with dotted lines. Peaks that were line-broadened beyond detection are indicated with grey circles. **(B)** Comparison of T2 relaxation times of free Fip1$_{226}$ (red), Fip1$_{226}$ + Yth1$_{ZF4}$ (green) and Fip1$_{226}$ + Pap1 (orange). Since residues in the Yth1-binding site are exchange broadened, only the first 170 residues on Fip1 were analyzed. Peaks that are significantly line broadened in Fip1$_{226}$ + Pap1 were not included for analysis.

**Supplemental Table S1: DNA oligonucleotides**

| Gene | Oligo | Sequence (5′-3′) |
|---|---|---|
| ***Sf*9 expression vectors** | | |
| Yth1$_{\Delta ZF45C}$ | Yth1-45CF | GGAATTCGGATCCCTCGAGATGAGCCTGATTCATCC |
| Yth1$_{\Delta ZF45C}$ | Yth1-45CR | GCCCCATCTAGAGGTACCTCATTACGGATCAATATGCAGATATTG |
| Yth1$_{\Delta ZF5C}$ | Yth1-5CF | GGAATTCGGATCCCTCGAGATGAGCCTGATTCATCC |
| Yth1$_{\Delta ZF5C}$ | Yth1-5CR | GCCCCATCTAGAGGTACCTCATTAATCTTTACCCAGAGGGC |
| Yth1$_{\Delta ZF4}$ | Yth1-del4F1 | GGAATTCGGATCCCTCGAGATGAGCCTGATTCATCC |
| Yth1$_{\Delta ZF4}$ | Yth1-del4R1 | ATGTTCCATATCACATTCCGGATCAATATGCAGATATT |
| Yth1$_{\Delta ZF4}$ | Yth1-del4F2 | AATATCTGCATATTGATCCGGAATGTGATATGGAACAT |
| Yth1$_{\Delta ZF4}$ | Yth1-del4R2 | ATTATTAACGGTGAAGTTTAATAATGAGGTACCTCTAGATGGGGC |
| Fip1$_{\Delta 1\text{-}60}$ | Fip1-d60F | GGAATTCGGATCCCTCGAGATGGATGATCGTAGTGATGAAGA |
| Fip1$_{\Delta 1\text{-}60}$ | Fip1-d60R | GCCCCATCTAGAGGTACCTCATTATTATTTGCTATTCTGGTTCTGATTCTG |
| Fip1$_{\Delta 110\text{-}180}$ | Fip1-dLCR1 | GGAATTCGGATCCCTCGAGATGAGCAGCAGCGAAGATG |
| Fip1$_{\Delta 110\text{-}180}$ | Fip1-dLCR2 | CGGTTTTTCTTTCAGAACTTCCGGATCGCTGCTGGTTGCTGCGGTGCTA |
| Fip1$_{\Delta 110\text{-}180}$ | Fip1-dLCR3 | TAGCACCGCAGCAACCAGCAGCGATCCGGAAGTTCTGAAAGAAAAACCG |
| Fip1$_{\Delta 110\text{-}180}$ | Fip1-dLCR4 | GCCCCATCTAGAGGTACCTCATTATTATTTGCTATTCTGGTTCTGATTCTG |
| Fip1$_{226}$ | Fip1-226F | GGAATTCGGATCCCTCGAGATGAGCAGCAGCGAAGATGAA |
| Fip1$_{226}$ | Fip1-226R | GCCCCATCTAGAGGTACCTCATTAATTGTAATCCTGCTGCAG |
| ***E. coli* expression vectors** | | |
| Fip1$_{226}$ | Fip1-226EcF | GGAATTCCATATGAGCAGCAGCGAAGATGAA |
| Fip1$_{226}$ | Fip1-226EcR | CCCAAAGCTTTTATTAATTGTAATCCTGCTGCAG |
| Yth1$_{\Delta ZF45C}$ | Yth1-45CEcF | ATTAAGGATCCATGGCAAGCAAAATTCCGAAA |
| Yth1$_{\Delta ZF45C}$ | Yth1-45CEcR | ATAATGAATTCTTATTATTAAACTTCACCGTTAATAATGGCG |
| Yth1$_{ZF4}$ | Yth1-ZnF4F | ATTAAGGATCCCCGGATTGTCAATAT |
| Yth1$_{ZF4}$ | Yth1-ZnF4R | ATAATGAATTCTTATTAATCTTTACCCAGAGGG |

**Supplemental Table S2: Buffers**

| Name | Composition |
|---|---|
| CPF lysis buffer | 200 mM HEPES pH 8, 200 mM KCl, 0.5 mM Mg(OAc)$_2$, 1 mM TCEP, 10% w/v glycerol, 2 μg/ml DNaseI (add fresh), 2 μg/ml RNaseA (add fresh) and protease inhibitor tables (14 tablets per 120 ml of buffer) (Roche) |
| CPF wash buffer | 50 mM HEPES pH 8, 150 mM KCl, 0.5 mM Mg(OAc)$_2$, 1mM TCEP |
| CPF strep elution buffer | CPF wash buffer supplemented with 1.2 mg/ml desthiobiotin |
| CFIA lysis buffer | 50 mM HEPES pH 7.9, 250 mM NaCl, 0.5 mM TCEP, 5% w/v glycerol, 2 μg/ml RNaseA (add fresh) and protease inhibitor tables (10 tablets per 120 ml of buffer) (Roche) |
| CFIA wash buffer | 20 mM HEPES pH 7.9, 250 mM NaCl, 0.5 mM TCEP |
| CFIA strep elution buffer | CFIA wash buffer supplemented with 1.2 mg/ml desthiobiotin |
| Pulldown lysis buffer | 100 mM HEPES pH 8, 300 mM NaCl, 5% glycerol, 1 mM TCEP |
| Pulldown wash buffer | 100 mM HEPES pH 8, 300 mM NaCl, 1mM TCEP |
| Pulldown elution buffer | 100 mM HEPES pH 8, 300 mM NaCl, 1mM TCEP, 6 mM desthiobiotin |
| Buffer A (Yth1) | 50 mM HEPES pH 7.4, 150 mM NaCl |
| Buffer B (Fip1) | 50 mM HEPES pH 7.4, 500 mM NaCl |
| Pap1 lysis buffer | 50 mM HEPES pH 8.0, 1 M NaCl, 20 mM imidazole, 5% w/v glycerol, 1 mM TCEP, 2 μg/ml Dnase I, 2 μg/ml Rnase A and protease inhibitor mixture (Roche) |
| Pap1 buffer | 50 mM HEPES pH 8.0, 1 M NaCl, 0.5 mM TCEP |
